# Identification of the Inducible HIV reservoir in Tonsillar, Intestinal and Cervical Tissue Models of HIV Latency

**DOI:** 10.1101/2025.02.10.637439

**Authors:** A. Gallego-Cortés, N. Sánchez-Gaona, C. Mancebo-Pérez, O. Ruiz-Isant, A. Benítez-Martínez, S. Landolfi, J. Castellví, F. Pumarola, N. Ortiz, I. Llano, J. Lorente, L. Mañalich-Barrachina, JG. Prado, E. Martín-Gayo, V. Falcó, M. Genescà, MJ. Buzon

## Abstract

HIV persists in diverse tissues, with distinct cellular reservoirs presenting a major barrier to a cure and requiring targeted therapeutic strategies to address this heterogeneity. Here, we developed tissue models of HIV latency using human tonsillar, intestinal and cervicovaginal tissues. These models revealed differential HIV infection across CD4^+^ T cell subpopulations, with ART partially restoring CD4^+^ T cells and reducing intact HIV DNA. T follicular helper cells (T_FH CD69+ CCR7-_) were the primary inducible reservoir in tonsils, while tissue-resident memory cells (T_RM CD69+ CD49a+_) dominated in the intestine. Identification of markers for inducible reservoirs revealed that CD69, CD45RO, and PD-1 were shared across tissues, while CXCR5 in the tonsils and CD49a in the intestine served as tissue-specific markers. Furthermore, using different latency reversal agents (LRAs) we found that Histone Deacetylase Inhibitors (HDACis) failed to induce HIV in any tissue, the SMAC mimetic AZD5582 was effective only in a resident-memory CD4^+^ T cell subpopulation in the intestine, and IL15 exhibited the broadest reactivation potential across tissues and CD4^+^ T subsets. These models recapitulate key aspects of HIV infection providing insights into the inducible reservoir’s composition in different tissues and informing strategies for its elimination.

## Introduction

Despite significant progress in antiretroviral therapy (ART), HIV infection is still an incurable disease^1,2^. While ART effectively suppresses HIV viremia in the bloodstream of people with HIV (PWH), it cannot completely eradicate the virus from the body due to the establishment of viral reservoirs. These reservoirs are mainly comprised of a pool of long-lived HIV-infected cells generated early after infection^3,4^ and upon ART initiation^5^, which harbour integrated, intact HIV DNA capable of producing replicative viruses and largely evading the immune system^6–8^. HIV reservoirs are distributed across diverse cell subtypes and tissue compartments^9–17^, representing the primary barrier to curing HIV. Thus, identifying and understanding the distinctive characteristics of HIV reservoirs’ cells is essential for developing strategies aimed at eliminating HIV from the body.

HIV infects multiple immune cell populations^18,19^, but memory CD4^+^ T cells are recognized as the primary contributors to the long-lived viral reservoir^20–22^. Specifically, central memory (T_CM_), transitional memory (T_TM_) and effector memory (T_EM_) CD4^+^ T cells are identified as major cellular reservoirs of latent HIV in peripheral blood of PWH on ART^21–24^. The pivotal role of less differentiated cell populations in HIV persistence has also been described^25,26^. While cellular reservoirs have been extensively studied in the bloodstream of PWH, the majority of CD4^+^ T cells, and consequently viral reservoirs, reside in tissues. Therefore, it is crucial to investigate HIV persistence within tissue compartments.

HIV has been detected in most tissues throughout the human body, including critical sanctuary sites such as lymphoid tissues (LT), the gastrointestinal tract, the central nervous system, and the reproductive tract ^17,27,28^. These anatomical compartments may sustain low levels of viral production due to their immune-privileged environment^29–32^ or reduced penetration of antiretroviral drugs^33–35^. Furthermore, clonal expansion of latently infected CD4^+^ T cells represents another mechanism contributing to HIV persistence in these tissues^36–38^. Compelling evidence indicate that lymph nodes (LN), gut mucosa and gut-associated lymphoid tissue (GALT) serve as major reservoirs for HIV in PWH on ART^39–41^. Moreover, a higher viral burden has been observed in the gastrointestinal and cervical tissues compared to peripheral blood in PWH on suppressive treatment^9,41,42^. In these tissue reservoirs, a significant proportion of the HIV resides within CD4^+^ T cells, with varying phenotype and proportion across tissues, adding to the complexity of HIV reservoirs. Therefore, it is imperative to identify the specific CD4^+^ T cell reservoirs within these tissues and develop strategies to purge them.

Certain CD4^+^ T cell populations, such as follicular helper T cells (T_FH_) in LN and type 17 helper T cells (T_H_17) in the gut, are pivotal in sustaining persistent HIV infection within these tissues^43–45^. Moreover, tissue-resident memory T cells (T_RM_) have been recently identified as a critical cellular reservoirs in the cervix and LN^9,46^. T_RM_ cells are a non-circulating memory subpopulation found in multiple tissues which mediate rapid protective responses to site-specific infections^47^. The presence of T_RM_ within distinct locations drives the development of T_RM_ cells functionally and phenotypically diverse, yet all T_RM_ cells share a core gene and protein profile^47–49^. They are characterized by the expression of CD69, which confers tissue retention properties^47,50^. T_RM_ can also express the adhesion molecules integrin αE (CD103) and integrin α1 (CD49a), the immune checkpoint receptor PD-1 and the IL7 receptor-alpha chain (CD127)^47,51,52^. Notably, T_RM_ are highly susceptible to HIV infection and possess enhanced proliferative capacity and longevity, making them potential targets for HIV persistence^9,47,53–56^. Moreover, the expression of PD-1 and CD127 on CD4^+^ T cells has been associated with a phenotype that predominantly supports latent HIV infection^13,14^. Our current understanding of how T_RM_ cells contribute to tissue reservoirs is limited. Considering their widespread distribution and abundance throughout the body, further research into the role of T_RM_ cells in HIV persistence is essential.

The diverse array of cellular reservoirs within tissues underscores the need to develop curative strategies capable of effectively targeting a wide range of cell populations to eradicate HIV. Recent efforts to eliminate the latent reservoir have focused on the “shock and kill” strategy. This approach uses latency reversal agents (LRAs) to reactivate HIV from latently infected cells, which are then cleared by virus-induced cytopathic effects or the immune system^57,58^. The effect of LRAs on circulating cellular reservoirs have been extensively studied^22,59–63;^ however, information regarding their impact on HIV reservoirs within tissues is scarce. Differential responses to current LRAs have been observed among the various CD4^+^ T cell subsets from the blood^22,59,61^, suggesting that cell populations within tissue compartments may also exhibit varying responses. Characterizing the susceptibility of distinct tissue CD4^+^ T cell subpopulations to pharmacological HIV reactivation is crucial for designing more effective therapies aimed at reducing HIV reservoirs. Nevertheless, the limited and challenging accessibility to tissue samples from PWH hinders this study.

In our study, we established models of latent HIV infection using human tonsillar, intestinal and cervicovaginal explants. We aimed to characterize productive and persistent viral infection, identify inducible HIV reservoirs, and evaluate the efficacy of various LRAs in these tissues. Our results showed a differential distribution of HIV infection among CD4^+^ T subpopulation across tissues. We also demonstrated the presence of inducible HIV reservoirs in our tissue models, primarily localized within T_FH_ cells in the tonsils and T_RM_ subsets in the intestinal tissue. Importantly, our study revealed distinct reactivation patterns of several LRAs, not only between tissues but also across different CD4^+^ T cell populations in these tissues, highlighting the tissue-specific and cell-specific nature of both HIV reservoir establishment and viral reactivation. These findings underscore the importance of developing therapeutic strategies with a broader spectrum of activity to target the different cellular reservoirs within anatomical compartments.

## Results

### Development of explant models for the study of HIV persistence in tissues

We initially worked on the establishment of three different explant models to characterize persistent HIV infection in human tissues. Tonsillar, intestinal and cervicovaginal explants were obtained from routine surgeries of uninfected donors and dissected into small blocks. Tissue blocks were cultured and HIV-infected as described in the schematic representation in **Figure 1A**. First, we identified the optimal conditions for establishing HIV infection in the explant models. To achieve this, we monitored the progression of infection by assessing intracellular p24 levels in CD4^+^ T cells throughout both productive and treated viral infection. Longitudinal analyses were performed for the tonsillar and intestinal models but were not feasible for the cervicovaginal model due to the limited size of the explants. A robust and consistent HIV infection was evident by day 5 in the tonsillar tissue (median_TO_=1.1% p24^+^ in CD4^+^ T cells, p_TO_<0.0001) and by day 6 in the intestinal and cervical tissues (median_GUT_= 3.6% p24^+^ in CD4^+^ T cells, p_GUT_<0.0001; median_CVX_= 0.75% p24^+^ in CD4^+^ T cells, p_CVX_=0.06) (**Figure 1B-C**). Intestinal explants consistently exhibited higher levels of HIV infection compared to tonsillar and cervicovaginal explants and ART administration effectively and significantly reduced HIV infection in all tissues within two to three days (median decrease in p24 expression: TO= 77%, GUT= 81% and CVX= 84%) (**Figure 1C-D**). Notably, the production of p24 was not entirely abrogated by ART (median_TO_= 0.5%, median_GUT_=2.3% and median_CVX_=0.05%) (**Figure 1D**). Additionally, we analysed longitudinal changes in key immune populations in the tonsillar and intestinal models, including CD4^+^ T and CD8^+^ T cells; NK cells (total CD56^+^ NK cells, as well as CD56^+^ CD16^+^ and CD56^+^ CD16 subsets) and B cells. An example of the gating strategy used for these analyses is shown in **Supplementary Figure 1A-B.** Overall, HIV infection did not cause significant alterations in the proportion of CD8^+^ T cells, NK cells and B cells (**Supplementary Figure 2**).

**Figure 1.**
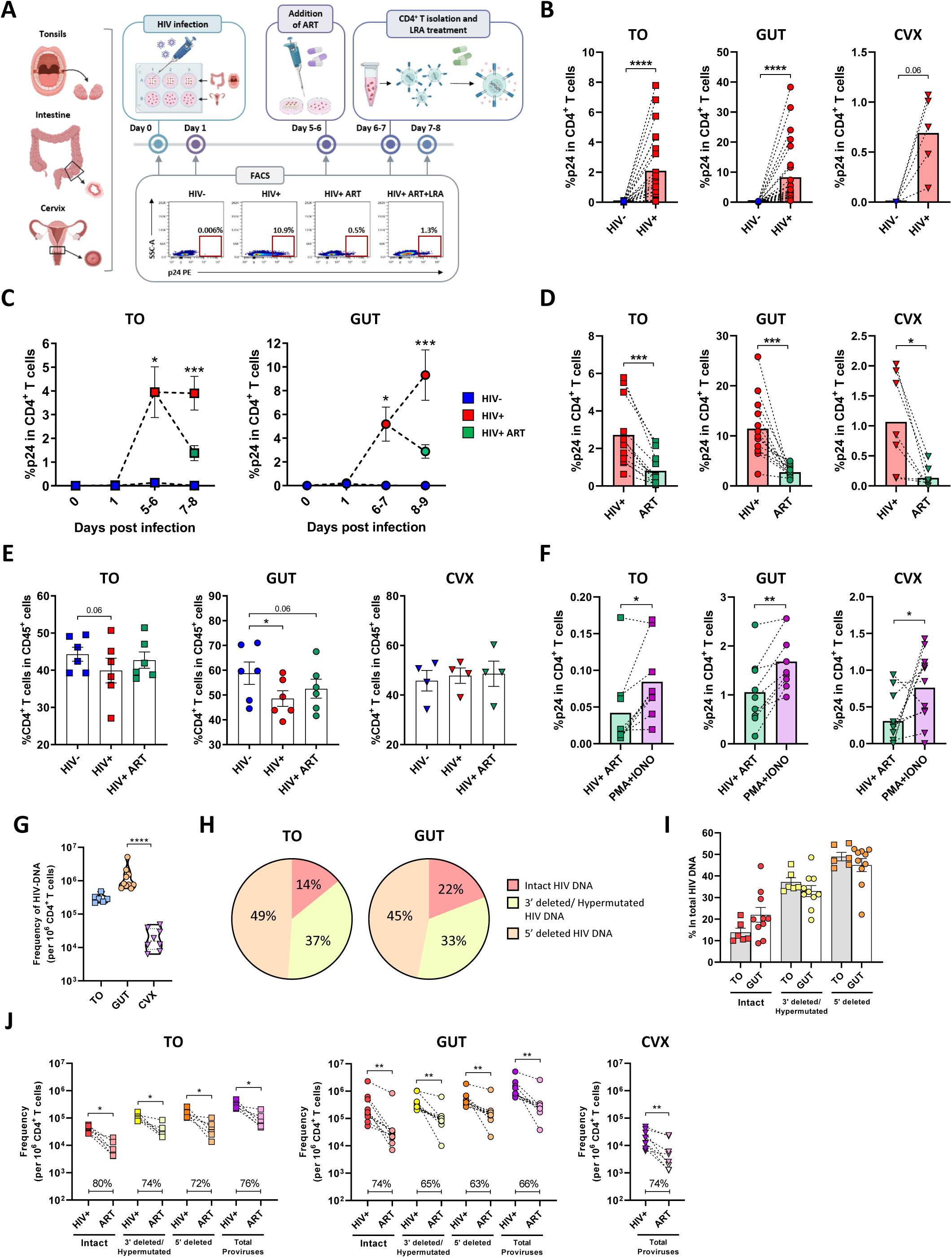
Tonsillar, intestinal and cervical models of latent HIV infection. (A) Schematic representation of the experimental workflow for studying productive and latent HIV infection in human tonsillar (TO), intestinal (GUT) and cervicovaginal (CVX) tissues. Tissue resections are dissected into small blocks, either placed on sponges or immersed in culture media, and subsequently infected with HIV_BAL_. After 5-6 days of infection, ART is added for 2 days, and CD4^+^ T cells are then isolated and cultured overnight with an LRA. Immune population changes and HIV infection levels (p24) are analysed at various time points. (B) Productive HIV infection in tonsillar, intestinal and cervical tissues. Graphs display p24 expression in CD4^+^ T cells from uninfected and infected tonsillar tissues (n=23) at day 5, and intestinal (n=26) and cervical tissues (n=5) at day 6. (C) Longitudinal monitoring of HIV infection in tonsillar and intestinal models. Graphs illustrate p24 levels in CD4^+^ T cells under uninfected, infected, and infected and ART- treated conditions in tonsillar (n=6) and intestinal (n=6) tissues over 8-9 days. Mean values and standard error of means (SEM) are shown. (D) Effect of ART administration on HIV infection. Graphs show p24 levels in CD4^+^ T cells from infected and infected and ART-treated tonsillar (n=12) tissues at days 7-8, and intestinal (n=13) and cervical (n=7) tissues at days 8-9. (E) Impact of HIV infection and ART on CD4^+^ T cells. Frequency of CD4^+^ T cells within CD45^+^ live cells under uninfected, infected, and infected and ART-treated conditions in all tissues at days 7-9. Mean values and SEM are depicted. (F) Viral reactivation of the inducible reservoir in tissue models. Tissues were infected as outlined in panel A. CD4^+^ T cells were isolated, and viral reactivation was assessed by measuring p24 levels after pharmacological stimulation. Phorbol 12- myristate 13-acetate (PMA) and ionomycin were used as LRAs. The graphs display p24 levels in both unstimulated and reactivated CD4^+^ T cells from tonsillar (n=9) tissues at day 8, and from intestinal (n=9) and cervical tissues (n=11) at day 9. (G) HIV reservoir size in infected tissues. Violin plots displaying the size of the HIV reservoir, measured as copies of HIV DNA per million isolated infected CD4^+^ T cells, from tonsillar tissue at day 7, and from intestinal and cervicovaginal tissues at day 8. (H) Composition of HIV DNA forms in the tonsillar and intestinal models during productive infection. Pie charts depict the percentages of intact, hypermutated, and 5’ deleted forms of HIV DNA within infected tonsillar (n=6) and intestinal (n=10) tissues at days 7 and 8, respectively. Mean values are shown. (I) Differences in the percentages of intact, hypermutated, and 5’ deleted forms of HIV DNA in isolated CD4^+^ T cells from productively infected tonsillar and intestinal tissues at days 7 and 8, respectively. Mean values and SEM are represented. (J) Composition of HIV DNA forms in tissue models during productive infection and after ART. Tissues were infected and treated with ART as outlined in panel A. Graphs show the quantification of intact, hypermutated, and 5’ deleted forms, as well as total HIV DNA in CD4^+^ T cells from infected and ART-treated tonsillar (n=6) and intestinal (n=10) tissues at days 7 and 8, respectively; and total HIV DNA in cervical tissues (n=8) at day 8. Medians of percentages of reduction for all forms of HIV DNA are indicated. Empty dots represent values below the limit of detection. Statistical comparisons were made using the Wilcoxon test in panels B, C, D, F, and J; the Friedman test in C and E; the Kruskal Wallis test in G; and the Mann-Whitney test in I. Significance levels are denoted as *p < 0.05; **p < 0.01; ***p < 0.001; ****p < 0.0001.

However, viral infection reduced the percentage of CD4^+^ T cells in both tissues, which was partially reverted after ART introduction (**Figure 1E, Supplementary Figure 2**). We also examined the impact of HIV infection and ART administration on CD4^+^ T cells within cervical tissue; however, no significant changes were observed, potentially due to fluctuations in the proportions of other cellular populations that may have overshadowed these effects (**Figure 1E**). Next, we evaluated the presence of inducible reservoirs by triggering viral reactivation in the tissue models. Pharmacological stimulation using PMA and Ionomycin on isolated ART-treated CD4^+^ T cells from all tissues resulted in a significant increase in intracellular p24 levels (median increase in p24 expression: TO= 69%, GUT= 35% and CVX= 44%), confirming the presence and reactivation of HIV reservoirs within tissues (**Figure 1F**). Interestingly, p24 expression under PMA/Ionomycin stimulation was higher in the cervix (median= 0.85%) compared to the tonsils (median= 0.07%) (**Figure 1F**), despite the cervix having lower levels of productive infection (**Figure 1B,D**). This discrepancy may be due to the smaller number of cells isolated from cervicovaginal tissue, which could overestimate its reactivation potential.

Finally, we analysed the size of the established HIV reservoirs within the tissues after 7- 8 days of productive infection. The Intact Proviral DNA Assay (IPDA) was performed on isolated CD4^+^ T cells from tonsillar and intestinal tissue blocks to quantify the different forms of HIV DNA (intact, 3’ deleted/hypermutated and 5’ deleted variants). Due to the limited number of CD4^+^ T cells isolated from cervicovaginal tissue, we measured only the amount of total HIV DNA using quantitative PCR (qPCR). Intestinal CD4^+^ T cells harboured the most HIV DNA, followed by tonsillar and cervicovaginal cells (**Figure 1G**), consistent with p24 expression levels (**Figure 1D**). When quantifying the distinct HIV DNA forms, we observed that 14% of the proviruses in tonsillar CD4^+^ T cells exhibited intact DNA, compared to 22% in CD4^+^ T cells from the intestine (**Figure 1H**). In both tissues, the 5’-deleted variant was the most prevalent form (mean_TO_= 49%, mean_GUT_= 45%), closely followed by the hypermutated variant (mean_TO_= 37%, mean_GUT_= 33%). Notably, there were no statistically significant differences in the proportions of the HIV DNA forms between the tissues (**Figure 1I**). After ART administration, we observed a significant decline in total HIV DNA levels across all tissues (**Figure 1J**). All forms of HIV DNA in tonsillar and intestinal tissues were reduced, with the decrease of the intact HIV DNA form being slightly more pronounced (**Figure 1J**).

Overall, we successfully established optimal models to study HIV persistence while on ART in tonsillar, intestinal and cervicovaginal tissues. These models recapitulate some of the major features of HIV infection, including the loss of CD4^+^ T cells after HIV infection, the protective role of ART against this decline, and the impact of ART on the HIV DNA. Furthermore, we demonstrated that these established models are responsive to pharmacological viral reactivation.

### Distinct representation of CD4^+^ T cell subpopulations across tissues and differential contribution to HIV infection in tonsillar and intestinal compartments

We then characterized the subset composition and phenotypic heterogeneity of the CD4^+^ T cell repertoires in tonsillar, intestinal and cervicovaginal tissues. CD4^+^ T cells isolated from the tissue blocks were analysed via flow cytometry using a comprehensive panel of markers for T cell differentiation, function and migration. Through unsupervised clustering based on the expression profiles of these markers (**Supplementary Figure 3A-B**), we identified 12 clusters of CD4^+^ T cells (**Figure 2A-B**). These CD4^+^ T subsets included naïve T cells (C01-02 T_NA_: CD45RO^-^ CCR7^+^), central memory T cells (C03, T_CM_: CD45RO^+^ CCR7^+^), effector memory T cells (C04, T_EM_: CD45RO^+^ CCR7^-^) and terminal effector memory T cells (C05-06, T_TEM_: CD45RO^-^ CCR7^-^). T_NA_ and T_TEM_ populations were further differentiated based on the expression of the activation marker CD69: clusters C01 and C05 represented cells in a resting state (CD69^-^), while cluster C02 and C06 showed stimulated phenotypes (CD69^+^). Moreover, we identified follicular helper T cells (C07-09), characterized by the expression of the chemokine receptor CXCR5, which facilitates their proximity to B cells within lymph nodes, and PD-1^64^ (T_FH_: CXCR5^+^ PD- 1^+^). These T_FH_ cells were further classified based on the differential expression of CCR7, which, together with CXCR5, is crucial for their positioning within lymph nodes and maturation^65,66^. Additionally, T_FH_ cells were categorized based on the presence or absence of CD69, a receptor also associated with tissue retention^47^, distinguishing between resident (CD69^+^) and non-resident T_FH_ cells (CD69^-^)^67^. Lastly, we identified three clusters of resident memory T cells, characterized by the essential expression of CD69 and the downregulation of CCR7, to ensure their retention within non-lymphoid tissues, as well as a memory phenotype (C10-12, T_RM_: CD45RO^+^ CCR7^-^ CD69^+^). These T_RM_ clusters differed in their expression of the adhesion molecules CD103 and CD49a, which determine their localization in tissue; as well as PD-1 expression^47^. The expression of the homeostatic cytokine IL7 receptor CD127, commonly present in long-living memory T cells^68^, was analysed in the identified subsets and found to be higher in T_RM_ clusters (C11 and C12), T_FH_ cells (C07 and C08) and in the T_CM_ subset (C03) (**Figure 2B, Supplementary Figure 3A**). Importantly, tissue memory CD4^+^ T cells expressing CD127 have previously been shown to contain latent HIV and be prone to viral reactivation^14^. Additionally, we assessed the expression of KLRG1, a marker associated with a more differentiated CD4^+^ T cell profile^69,70^ which has also been identified as an immune inhibitory checkpoint receptor^71^. KLRG1 is expressed on CD4^+^ T cells harbouring inducible HIV genomes^71^ and clonally expanded proviruses^46^. We found that the T_RM_ cluster C11 exhibited slightly higher KLRG1 expression (**Figure 2B, Supplementary Figure 3A**).

**Figure 2.**
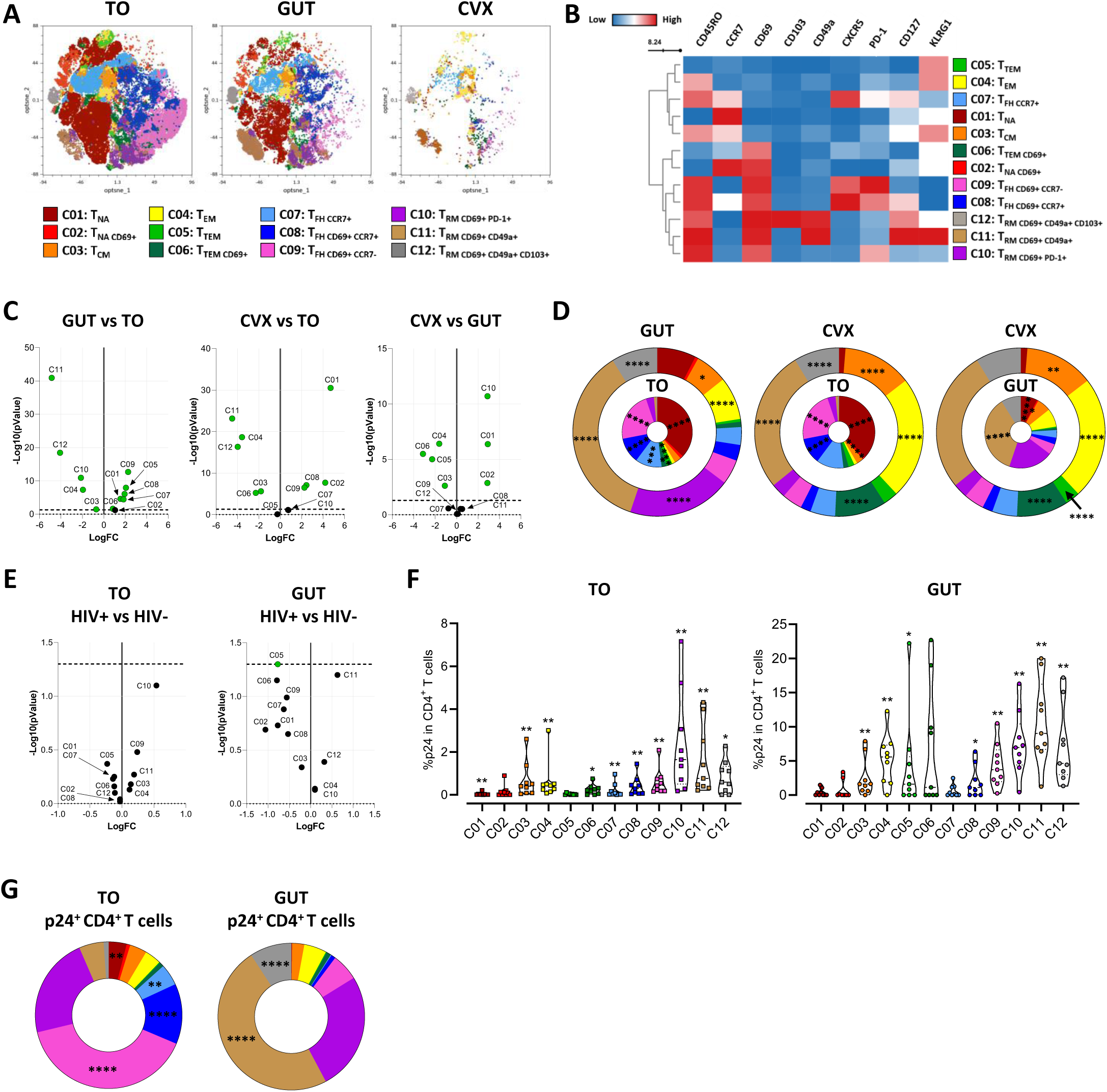
Distinct CD4^+^ T cell compartments in the tissue models and differential distribution of HIV infection in the tonsillar and intestinal tissues. (A) Opt-SNE plots displaying the distribution of the twelve cell clusters identified based on the expression of CD45RO, CCR7, CD69, CD103, CD49a, CXCR5, PD-1, CD127 and KLRG1 within isolated CD4^+^ T cells from uninfected tonsils at day 8, and intestines and cervix at day 9. (B) Heatmap illustrating differences in the Mean Fluorescence Intensity (MFI) of the studied markers across the twelve identified clusters. (C) Volcano plots showing the differences in the proportion of clusters between the tissues. The edgeR statistical method is employed, with green dots indicating statistically significant differences. (D) Pie charts representing the median percentages of each cluster within the total CD4^+^ T cell pool from uninfected tonsillar, intestinal and cervical tissues. Pairwise comparisons are made between the intestinal (outer circle) and tonsillar (inner circle), cervical (outer circle) and tonsillar (inner circle), and cervical (outer circle) and intestinal (inner circle) tissues. Statistical values are derived from the volcano plots in panel C, with significance levels indicated (*p < 0.05; **p < 0.01; ***p < 0.001; ****p < 0.0001). (E) Volcano plots showing the statistically significant differences in the proportion of clusters between the uninfected and infected conditions in both tissues. The edgeR statistical method is employed, with green dots indicating statistically significant differences. (F) Levels of HIV infection in CD4^+^ T cell clusters. Graphs illustrating p24 expression within each cluster of infected CD4^+^ T cells from tonsillar (n=9, day 8) and intestinal (n=9, day 9) tissues. Medians and quartiles are represented, and Wilcoxon test results comparing uninfected and infected conditions are shown (*p < 0.05; **p < 0.01). (G) Pie charts depicting the contribution of each cluster to the total HIV infection levels in the pool of infected CD4^+^ T cells from both tonsillar and intestinal tissues. Percentages are calculated based on the count of p24^+^ cells in each cluster relative to the total number of p24^+^ CD4^+^ T cells. Medians are shown, and statistical comparisons between tissue types were performed using the Mann-Whitney test (*p < 0.05; **p < 0.01; ***p < 0.001; ****p < 0.0001).

Next, we compared the proportions of the identified clusters across tissues, revealing significant inter-tissue differences in the distribution of the majority of the subsets (**Figure 2C-D**). The tonsillar tissue exhibited a high abundance of less differentiated subsets and T_FH_ cells, consistent with existing literature^72–76^. Specifically, we observed a greater representation of clusters C01:T_NA_ and C09:T_FH CD69+ CCR7-_, comprising 37% and 23% of the pool of CD4^+^ T cells, respectively (**Figure 2D**). Conversely, T_RM_ clusters predominated in the intestinal tissue. In particular, clusters C10:T_RM CD69+ PD-1+_ and C11:T_RM CD69+ CD49a+_ were the most abundant, accounting for 20% and 36% of the total CD4^+^ T cells, respectively (**Figure 2D**). This prevalence of T_RM_ cells within the human intestinal CD4^+^ T cell compartment has been previously documented^54,77^. Similarly, in cervicovaginal tissue, cluster C11:T_RM CD69+ CD49a+_ was the most abundant cluster, comprising 27% of total CD4^+^ T cells, followed closely by cluster C04:T_EM_, which represented 24% of the pool (**Figure 2D**). These findings align with prior reports describing a predominance of T_EM_ and T_RM_ CD4^+^ T cells in the cervical immune environment ^9,78,79^.

We also evaluated HIV infection among all the identified CD4^+^ T subsets from the tonsils and the intestine. This analysis could not be performed for the cervicovaginal tissue due to the limited amount of CD4^+^ T cells isolated. Firstly, we assessed the impact of productive infection in the abundance of the different clusters (**Supplementary Figure 4A-B**). Despite the ability of HIV to modify the phenotype of CD4+ T cells^80–82^, no novel clusters emerged in the HIV-infected conditions and we only observed a slight yet significant difference in the proportion of cluster C05:T_TEM_ in the intestine (p= 0.049) (**Figure 2E**). The proportions of the other clusters remained consistent in both tissues, indicating that the analysed phenotypes were not significantly altered. We then quantified HIV infection across the distinct CD4^+^ T clusters by measuring the expression of the p24 viral protein. As expected, infection levels were higher in the intestinal compared to the tonsillar CD4^+^ T cells (**Figure 2F**). A significant HIV infection was observed across most subpopulations in tonsillar tissues, with notably higher infection levels detected in the T_RM_ clusters, particularly in C10:T_RM CD69+ PD-1+_ (**Figure 2F**). In the intestine, viral infection predominantly occurred within the T_RM_ clusters C10:T_RM CD69+ PD-1+_, C11:T_RM CD69+ CD49a+_ and C12:T_RM CD69+ CD49a+ CD103+_. Comparable levels of infection were detected in the subset C04:T_EM_, while lower yet significant levels of viral infection were observed in the clusters C03:T_CM_, C05:T_TEM_, C08:T_FH CD69+ CCR7+_ and C09:T_FH CD69+ CCR7-_ (**Figure 2F**). Interestingly, the subpopulation with the highest CD127 levels and slight KLRG1 expression (C11:T_RM CD69+ CD49a+_) was among the most significantly infected clusters in both tissues (**Figure 2F**). Ultimately, we determined the contribution of each cluster to the overall infection within the total population of CD4^+^ T cells (**Figure 2G**). The tonsils exhibited a dispersed distribution of infection but it primarily localized in T_FH_ and T_RM_ subsets. Clusters C08:T_FH CD69+ CCR7+_, C09:T_FH CD69+ CCR7-_ and C10:T_RM CD69+ PD-1+_ made the greatest contributions to total levels of infection accounting for 13%, 40%, and 22% of the total p24^+^ CD4^+^ T cells, respectively (**Figure 2G, Supplementary Figure 4C**). Moreover, the contributions of clusters C01:T_NA_, C07:T_FH CCR7+_, C08:T_FH CD69+ CCR7+_ and C09:T_FH CD69+ CCR7-_ to the overall infection were significantly higher in the tonsils than in the intestine (**Figure 2G**). In contrast, infection concentrated in the T_RM_ clusters C10:T_RM CD69+ PD-1+_, C11:T_RM CD69+ CD49a+_ and C12:T_RM CD69+ CD49a+ CD103+_ in the intestine, accounting for 26%, 49%, and 9% of the total p24^+^ CD4^+^ T cells respectively, with the contribution of C11 and C12 significantly surpassing those in the tonsils (**Figure 2G, Supplementary Figure 4C**).

Overall, our findings unveiled notable differences in the composition of the CD4^+^ T cell compartments across the studied tissues, with T_NA_ and T_FH_ cells predominating in the tonsils, while intestinal and cervical CD4^+^ T cells primarily exhibited a T_RM_ phenotype. The T_FH_ fraction in the tonsils harboured the greatest number of infected cells, while HIV infection was primarily localized within the T_RM_ subpopulations in the intestinal tissue. Moreover, the contribution of the CD4^+^ T subsets to the total pool of HIV-infected cells differed markedly between tonsillar and intestinal tissues.

### Tonsillar and intestinal inducible HIV reservoirs are constituted of distinct CD4^+^ T subpopulations

As HIV reservoirs are established promptly following primary initial infection and shortly after the initiation of ART^5^, we aimed to identify the specific CD4^+^ T cell clusters that constituted inducible HIV reservoirs within the pool of tonsillar and intestinal infected CD4^+^ T cells. To achieve this, tissue blocks were HIV-infected and treated with ART. Inducible viral reservoirs were identified after maximal cell activation. For that, we stimulated isolated ART-treated HIV-infected CD4^+^ T cells with PMA and Ionomycin (**Figure 1A**). First, we compared the proportions of CD4^+^ T cell clusters under unstimulated and PMA/Ionomycin-stimulated conditions, as treatment with these agents might induce phenotypic changes associated with T cell activation and differentiation^15,22,83–85^. Thus, as previously reported^22,84,86,87^, we anticipated an upregulation of CD69 and PD-1 expression, and a downregulation of CXCR5 and CD127. Besides significantly altering some of the CD69^-^ and non-T_RM_ CD69^+^ clusters (C01:T_NA_, C02:T_NA CD69+_, C03:T_CM_, C04:T_EM_, C05:T_TEM_, C06:T_TEM CD69+_ and C07:T_FH CCR7+_), the proportion of the T_RM_ subsets remained largely unchanged in both tissues (**Supplementary Figure 5A-B**). Additionally, T_FH_ cells expressing CD69 in the tonsils, which contributed the most to HIV infection, were similarly unaffected (**Supplementary Figure 5A**). This indicated that these cells retained their phenotype following stimulation with PMA and Ionomycin. One possible explanation is that CD69 expression was already elevated in these subsets and did not increase further following cell activation, as previously shown for T_RM_ in cervix^9^. Next, we measured p24 expression in the clusters following viral reactivation (**Figure 3A**). Five tonsillar and three intestinal clusters exhibited statistically significant viral reactivation, identifying them as inducible HIV reservoirs during ART. Specifically, cluster C10:T_RM CD69+ PD-1+_ showed significant reactivation in both tissues; clusters C03:T_CM_, C08:T_FH CD69+ CCR7+_ and C09:T_FH CD69+ CCR7-_ were exclusively reactivated in the tonsils; and clusters C11:T_RM CD69+ CD49a+_ and C12:T_RM CD69+ CD49a+ CD103+_ were selectively reactivated in the intestine (**Figure 3A**). Additionally, p24 levels were elevated in cluster C02:T_NA CD69+_ in the tonsil, despite not detecting significant infection in this cluster (**Figure 2F**). We believe that this might be due to the upregulation of CD69 in cells from cluster C01:T_NA_, which could transition to C02:T_NA CD69+_ upon activation.

**Figure 3.**
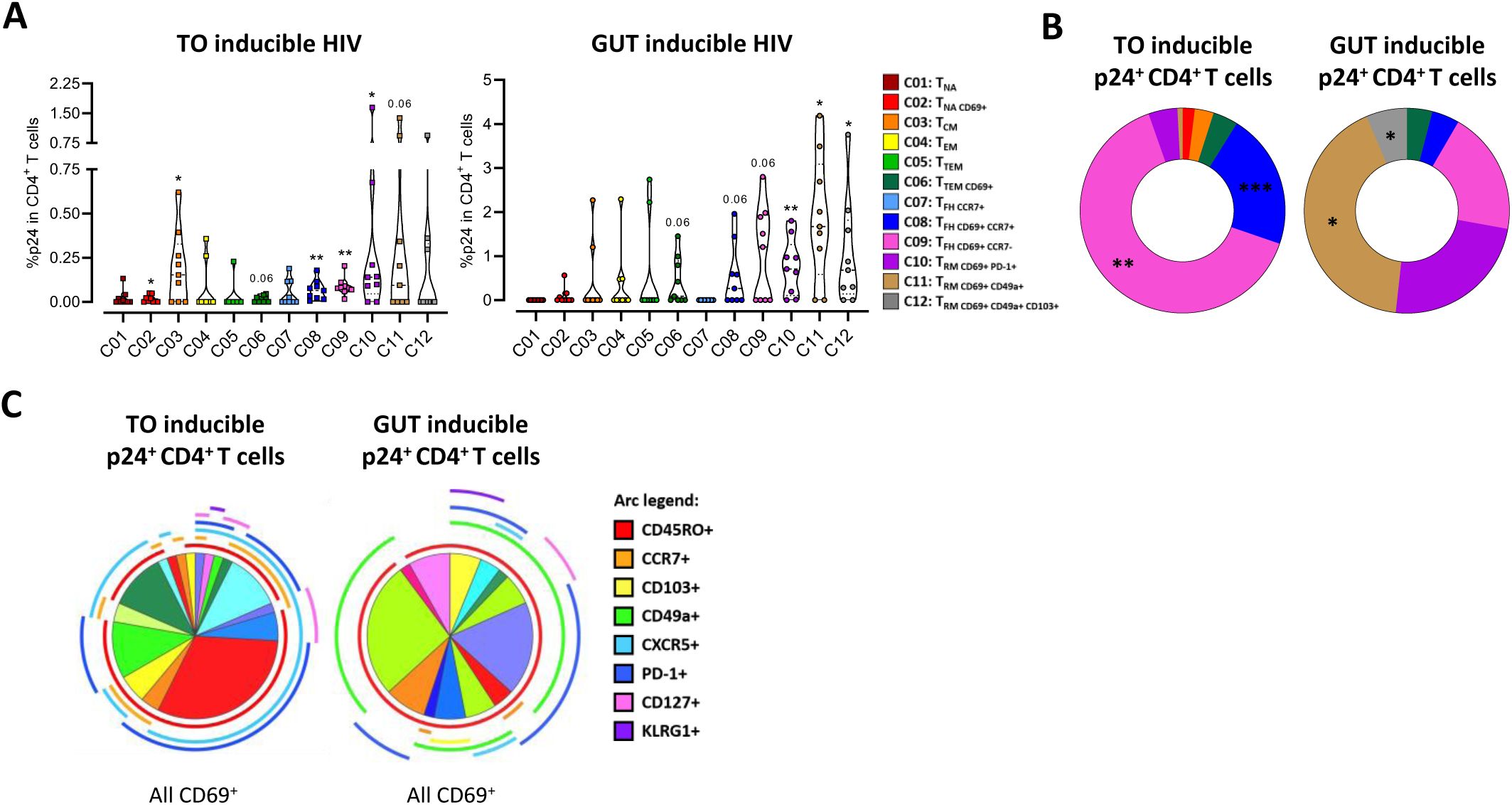
Tonsillar and intestinal inducible HIV reservoirs are constituted of distinct subpopulations of CD4^+^ T cells. (A) Inducible HIV reservoirs in CD4^+^ T cell clusters after viral reactivation with PMA and ionomycin. Graphs show the expression of p24 within each cluster of virally reactivated CD4^+^ T cells from tonsillar tissues (n=9) on day 8 and intestinal tissues (n=9) on day 9. Median values with quartiles are represented. Statistical comparisons of p24 levels between the unstimulated (ART) condition and the condition reactivated with PMA and ionomycin were conducted. (B) Pie charts illustrating the contribution of each cluster to the overall inducible reservoir in CD4^+^ T cells from tonsillar and intestinal tissues. Contribution is calculated as the count of p24^+^ cells within each cluster as a percentage of the total count of p24^+^ CD4^+^ T cells. Medians of these percentages are depicted, and statistical comparisons were performed. (C) Pie charts displaying the distribution of the different clusters derived from Boolean filters applied to phenotypic markers within the p24^+^ CD4^+^ T cell population. Arcs represent the expression of the phenotypic markers, each depicted in a distinct colour. All p24^+^ CD4^+^ T cells express CD69. Statistical comparisons were performed using the Wilcoxon test in (A); and the Mann-Whitney test in (B). Significance levels are denoted as *p < 0.05; **p < 0.01 and ***p < 0.001.

Subsequently, we analysed the contribution of each cluster to the overall inducible HIV reservoir in CD4^+^ T cells (**Figure 3B, Supplementary Figure 5C**). In the tonsils, cluster C09:T_FH CD69+ CCR7-_ was the primary contributor to total levels of inducible HIV, accounting for 64% of the p24^+^ CD4^+^ T cells. Reactivated cells from cluster C08:T_FH CD69+ CCR7+_ also comprised a significant portion, representing 21% of the total reactivated cells. Among intestinal cells, cluster C11:T_RM CD69+ CD49a+_ (the one expressing the highest levels of CD127 and KLRG1) predominated, representing 42% of virally reactivated cells. Cluster C10:T_RM CD69+ PD-1+_ also significantly contributed to the total inducible cell reservoir, accounting for 24%. Although cluster C09:T_FH CD69+ CCR7-_ did not consistently reactivate in all intestinal samples, it substantially contributed to the reactivated cell pool in those intestines where it did. Significant differences were observed in the contribution of certain clusters to reactivation between tissues. In particular, p24^+^ cells from clusters C08:T_FH CD69+ CCR7+_ and C09:T_FH CD69+ CCR7-_ were significantly more prevalent in the pool of reactivated tonsillar CD4^+^ T cells compared to their intestinal counterparts (**Figure 3B**). In contrast, clusters C11:T_RM CD69+ CD49a+_ and C12:T_RM CD69+ CD49a+ CD103+_ exhibited the opposite pattern (**Figure 3B**).

To further characterize the profile of reactivated cells and identify antigens that could define inducible HIV reservoirs in these tissues, we analysed the expression of the phenotypic markers within the pool of p24^+^ CD4^+^ T cells following PMA/Ionomycin stimulation (**Figure 3C**). In both tonsillar and intestinal tissues, reactivated cells consistently expressed CD69, and nearly all of them displayed the memory marker CD45RO. This may appear inconsistent with our previous observations, where T_CM_ cells, which lack CD69, reactivated significantly (**Figure 3A**); however, it is plausible that CD69 expression is induced in this cluster upon reactivation. Notably, the majority of the reactivated cells from the tonsils expressed CXCR5 and PD-1, whereas those from the intestine predominantly expressed CD49a and PD-1. CD49a and CD103 were absent in tonsillar reactivated cells, potentially due to the under-representation of T_RM_ cells within the tonsillar compartment (**Figure 2A-D**). These findings identify CD69, CD45RO, and PD-1 as shared markers of inducible reservoirs in both tonsillar and intestinal tissues. Additionally, CXCR5 emerges as predominantly associated with tonsillar reservoirs, whereas CD49a and, to a lesser extent, CD103 are specific to intestinal reservoirs.

Overall, while some clusters comprise potential HIV reservoirs in both tissues, there are unique tonsillar and intestinal CD4^+^ T cell reservoirs. In the tonsils, the primary reservoir is formed by T_FH_ cells, whereas T_RM_ populations constitute the largest inducible HIV reservoir in the intestinal tissue. Inducible HIV reservoirs within these tissue compartments shared CD69, CD45RO and PD-1 expression, while distinct markers, such as CXCR5 in the tonsils and CD49a in the intestine, delineated specific reservoir populations. Together, these results suggest both shared and tissue-restricted features of inducible HIV reservoirs across these compartments and identify potential marker candidates for characterizing inducible reservoirs in tonsils and intestinal tissues.

### Current LRAs demonstrated differing potency and breath of viral reactivation between tissue reservoirs

We aimed to evaluate the efficacy of different LRAs in inducing viral reactivation in the HIV-latency tissue models. The LRAs tested included the protein kinase C (PKC) agonist Ingenol (ING), histone deacetylase inhibitors (HDACis) Romidepsin (RMD) and Panobinostat (PNB), the SMAC mimetic AZD5582 (AZD), and interleukin IL15. Additionally, Ingenol and Romidepsin were evaluated in combination (I+R). As previously, the efficacy of the LRAs was determined by quantifying p24 expression after LRA-stimulation in isolated and ART-treated CD4^+^ T cells (**Figure 1F**). IL15 induced a potent HIV reactivation across all tissue reservoirs (**Figure 4A**). The combination of Ingenol and Romidepsin significantly reactivated HIV in tonsillar and intestinal CD4^+^ T cells, while Ingenol alone was sufficient to reactivate the virus in tonsillar and cervicovaginal reservoir cells (**Figure 4A**). AZD5582 demonstrated effectiveness exclusively in reactivating the intestinal reservoirs (**Figure 4A**). Subsequently, we compared the efficacy of the LRAs between tonsillar and intestinal tissues only, as the limited CD4^+^ T cell yield from cervicovaginal tissues precluded testing all LRAs in a single sample. A marked difference was observed in the efficacy of Ingenol between the intestinal and tonsillar reservoirs (**Figure 4B**).

**Figure 4.**
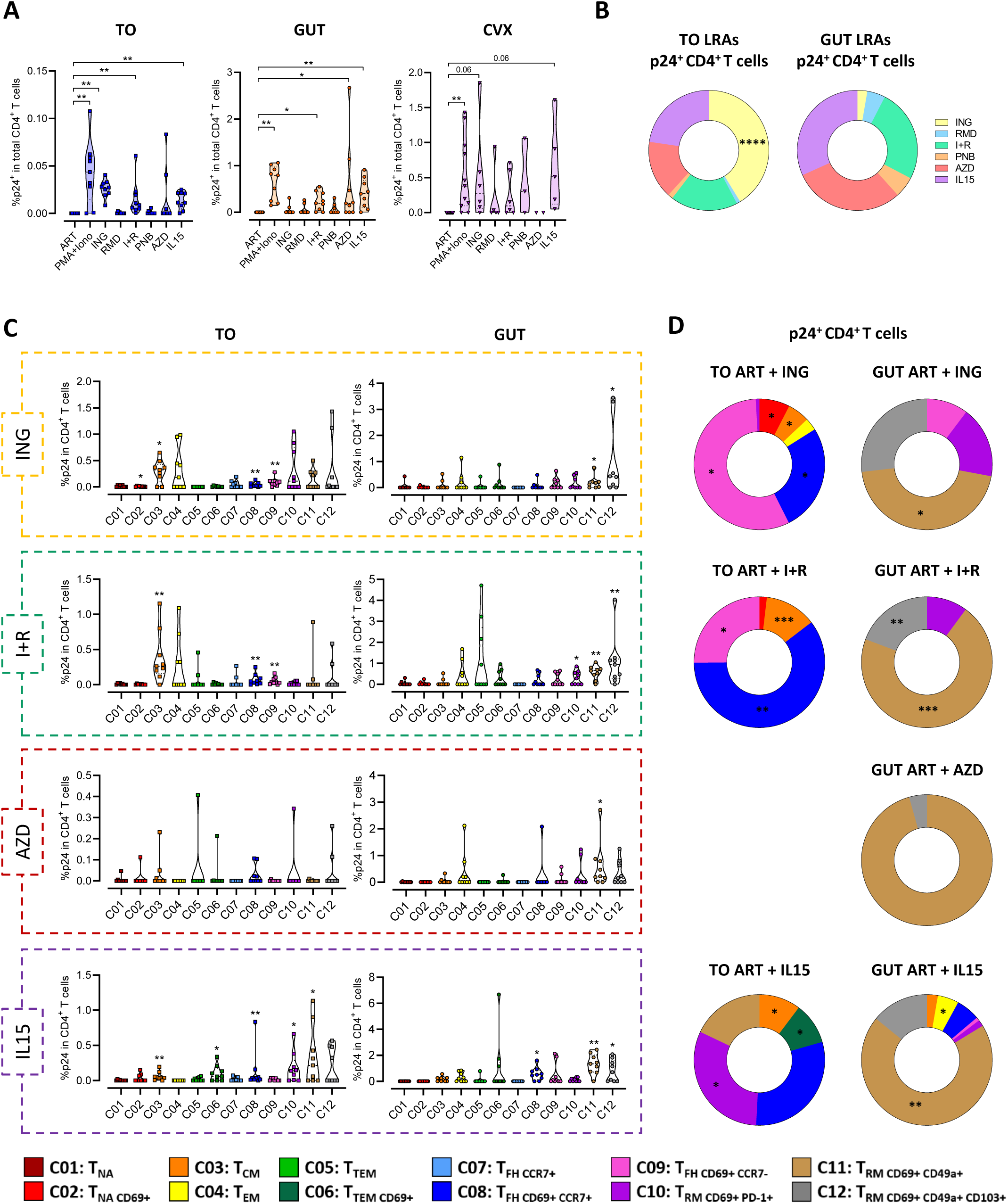
Differences in the efficacy of current LRAs in inducing viral reactivation of tissue HIV reservoirs. (A) Viral reactivation of the inducible reservoir of HIV-infected CD4^+^ T cells from tonsillar, intestinal and cervical tissues. Graphs display p24 expression within HIV-infected and ART-treated CD4^+^ T cells following stimulation with different LRAs: PMA and ionomycin (PMA+Iono), Ingenol (ING), Romidepsin (RMD), Ingenol and Romidepsin combined (I+R), Panobinostat (PNB), AZD5582 (AZD), and IL15. Median values with quartiles are represented. (B) Pies charts illustrating the contribution of the different LRAs to HIV reactivation in the total pool of virally reactivated CD4^+^ T cells from tonsillar and intestinal tissues. Means are depicted. (C) Effect of each LRA on viral reactivation in subpopulations of ART-treated CD4^+^ T cells from tonsil and intestine. Graphs display p24 expression within HIV-infected and ART-treated CD4^+^ T cells following stimulation with different LRAs: ING, I+R, AZD, and IL15. Median values with quartiles are represented. Statistical comparisons of p24 levels between the unstimulated (ART) condition and the condition reactivated with each LRA were conducted. (D) Pie charts illustrating the contribution of each cluster to overall HIV reactivation with ING, I+R, AZD, and IL15 in the total pool of virally reactivated CD4^+^ T cells from tonsillar and intestinal tissues. Contribution is calculated as the count of p24+ cells within each cluster, expressed as a percentage of the total p24^+^ CD4^+^ T cells. Medians of these percentages are depicted, and statistical comparisons between tissues were conducted. Statistical comparisons were performed using the Wilcoxon test in (A) and (C) and the Mann-Whitney test in (B) and (D) with significance levels denoted as *p < 0.05; **p < 0.01, ***p < 0.001; ****p < 0.0001.

Next, we evaluated the impact of these LRAs on the distinct subpopulations of CD4^+^ T cells, focusing on the clusters identified as potential HIV reservoirs in the tonsillar and intestinal tissues. We first examined whether the abundance of these CD4^+^ T subpopulations differed between the non-reactivated (ART) and LRAs-treated conditions (**Supplemental Figure 6A-B**). In the tonsils, the primary contributor to the inducible HIV reservoir, C09:T_FH CD69+ CCR7-_, remained consistently unaffected by all LRAs. The second major inducible cluster, C08:T_FH CD69+ CCR7+_, was reduced in the IL15 treatment condition and marginally increased with Ingenol, both alone and in combination with Romidepsin. This combination also minimally impacted the proportions of clusters C03:T_CM_ and C10:T_RM CD69+ PD-1+_ (**Supplementary** Figure 6A). In contrast, we observed consistent proportions of all clusters identified as inducible HIV reservoirs in the intestine (C10:T_RM CD69+ PD-1+_, C11:T_RM CD69+ CD49a+_ and C12:T_RM CD69+ CD49a+ CD103+_) before and after treatment with LRAs (**Supplementary** Figure 6B). Then, we measured p24 expression within each cluster after treatment with the different LRAs (**Figure 4C**). Ingenol alone and in combination with Romidepsin demonstrated a potent capacity to reactivate the tonsillar subsets C03:T_CM_, C08:T_FH CD69+ CCR7+_ and C09:T_FH CD69+ CCR7-_ and the intestinal clusters C11:T_RM CD69+ CD49a+_ and C12:T_RM CD69+ CD49a+ CD103+_ (**Figure 4C**). Moreover, Ingenol with Romidepsin successfully reactivated the intestinal reservoir C10:T_RM CD69+ PD-1+_. AZD5582 showed no effect in any of the tonsillar subsets but was highly effective in reactivating the intestinal T_RM_ reservoir C11:T_RM CD69+ CD49a+_. IL15 exhibited substantial reactivation capacity in the tonsillar reservoirs C03:T_CM_, C08:T_FH CD69+ CCR7+_ and C10:T_RM CD69+ PD-1+_ and the intestinal clusters C11:T_RM CD69+ CD49a+_ and C12:T_RM CD69+ CD49a+ CD103+_. Interestingly, while any other LRA failed to induce a significant reactivation in the tonsillar subsets C06:T_TEM CD69+_ and C11:T_RM CD69+ CD49a+_ and the intestinal cluster C08:T_FH CD69+ CCR7+_, IL15 could reactivate these clusters. Romidepsin and Panobinostat showed no reactivation potential in any tonsillar nor intestinal CD4^+^ T subpopulations (**Supplementary Figure 6C**). Importantly, despite being susceptible to HIV infection (**Figure 2F**), tonsillar clusters C04:T_EM_, C07:T_FH CCR7+_ and C12:T_RM CD69+ CD49a+ CD103+_ and intestinal clusters C03:T_CM_, C04:T_EM_, C05:T_TEM_ were not reactivated with any of the LRAs tested.

Finally, we assessed the contribution of each cluster to the total p24^+^ CD4^+^ T cells following stimulation with the LRAs (**Figure 4D**). In the tonsils, reactivated cells from clusters C08:T_FH CD69+ CCR7+_ and C09:T_FH CD69+ CCR7-_ were the most abundant in all LRA- stimulated conditions, with a minor yet significant contribution from cluster C03:T_CM._ Conversely, across most LRAs, the intestinal cluster C11:T_RM CD69+ CD49a+_ made the largest contribution to the total number of reactivated cells, followed by cluster C12:T_RM CD69+ CD49a+ CD103+_. Notably, ING and, to a lesser extent, IL15 were the most effective LRAs in the tonsils, while IL15 induced the broadest reactivation in the intestine (**Figure 4D**).

In summary, we observed that HIV reservoirs from the tonsils, intestine and cervix are more efficiently reactivated by Ingenol alone or in combination with Romidepsin, and IL15. However, at the level of CD4^+^ T cell subpopulations, LRAs induce reactivation in distinct subsets both across different tissues and within the same tissue. Specifically, while T_CM_ and T_FH_ cells in the tonsils and the T_RM_ fraction in the intestine were reactivated by most tested LRAs, the specific clusters affected differed between LRAs. Notably, IL15 was capable of reactivating populations in both tissues that remained unresponsive to other LRAs.

### Heterogeneous responses of tonsillar and intestinal CD4^+^ T cell subpopulations to LRAs

Finally, we evaluated the differential response of reservoir cells from the same clusters across tonsillar and intestinal tissues to the selected LRAs. Our previous data indicated that HIV predominantly persists in resident CD4^+^ T cells in both tissues, and that these subpopulations exhibited enhanced responsiveness to the tested LRAs. Nonetheless, inducible HIV reservoirs were also identified in the tonsillar clusters C03:T_CM_ and C06:T_TEM CD69+_ (**Figure 5A-B**). The reservoir in cluster C03:T_CM_ could be reactivated by Ingenol alone and in combination with Romidepsin, as well as with IL15. Remarkably, the combination treatment was most potent for cluster C03:T_CM_, achieving reactivation in nearly all tonsillar samples (**Figure 5A**). In contrast, reactivation of cluster C06:T_TEM CD69+_ was solely observed with IL15 (**Figure 5B**). In the tonsils, HIV reservoirs primarily localized within the resident T_FH_ fraction (C08:T_FH CD69+ CCR7+_ and C09:T_FH CD69+ CCR7-_), which notably exhibited the most robust reactivation response to the LRAs. However, differences were observed in the efficacy and extent of viral reactivation (**Figure 5C-D**). Both tonsillar clusters C08:T_FH CD69+ CCR7+_ and C09:T_FH CD69+ CCR7-_ responded to stimulation with Ingenol alone and in combination with Romidepsin, with subset C08:T_FH CD69+ CCR7+_ exhibiting a greater response to the combined treatment (**Figure 5C**); while cluster C09:T_FH CD69+ CCR7-_, to the single Ingenol (**Figure 5D**). Additionally, IL15 could significantly reactivate reservoir cells from cluster C08:T_FH CD69+ CCR7+_. Conversely, in the intestine, despite being significantly infected, these clusters were rarely virally reactivated by the LRAs. Only HIV present in cluster C08:T_FH CD69+ CCR7+_ was reactivated with IL15 (**Figure 5C**). As IL15 could reactivate HIV from cluster C08:T_FH CD69+ CCR7+_ in both tonsillar and intestinal tissues, we compared its potency between the tissues and observed that they exhibited similar reactivation efficiencies (**Supplementary Figure 7**). Ultimately, we evaluated the reactivation in the T_RM_ subsets, which comprise the majority of the HIV reservoir in the intestine but only a small proportion in the tonsils. The T_RM_ cluster C10:T_RM CD69+ PD-1+_ harboured HIV reservoirs in both tonsillar and intestinal tissues (**Figure 5E**), yet they were reactivated by distinct LRAs. Specifically, this reservoir was selectively reactivated by IL15 in the tonsils, and by the co-administration of Ingenol with Romidepsin in the intestine. The largest HIV reservoir in the intestinal tissue consisted of infected cells from cluster C11:T_RM CD69+ CD49a+_, which exhibit the highest levels of CD127 and slight KLRG1 expression. These cells could be reactivated by nearly all tested LRAs except for Romidepsin alone and Panobinostat (**Figure 5F**). Notably, this was the only cluster responsive to reactivation with AZD5582. The most potent reactivation was achieved with IL15, which also successfully reactivated the same cluster in the tonsils (**Figure 5F**). Moreover, the efficacy of IL15 was comparable in both the tonsillar and intestinal clusters C11:T_RM CD69+ CD49a+_ (**Supplementary Figure 7**). Finally, we analysed the response of cluster C12:T_RM CD69+ CD49a+ CD103+_ to the tested LRAs (**Figure 5G**). We observed that an inducible HIV reservoir was present exclusively in the intestinal cluster. Cells from this subset were successfully reactivated using Ingenol, both as a single treatment and in combination with Romidepsin, as well as with IL15. The combination of Ingenol and Romidepsin proved to be the most effective reactivation strategy for this subset.

**Figure 5.**
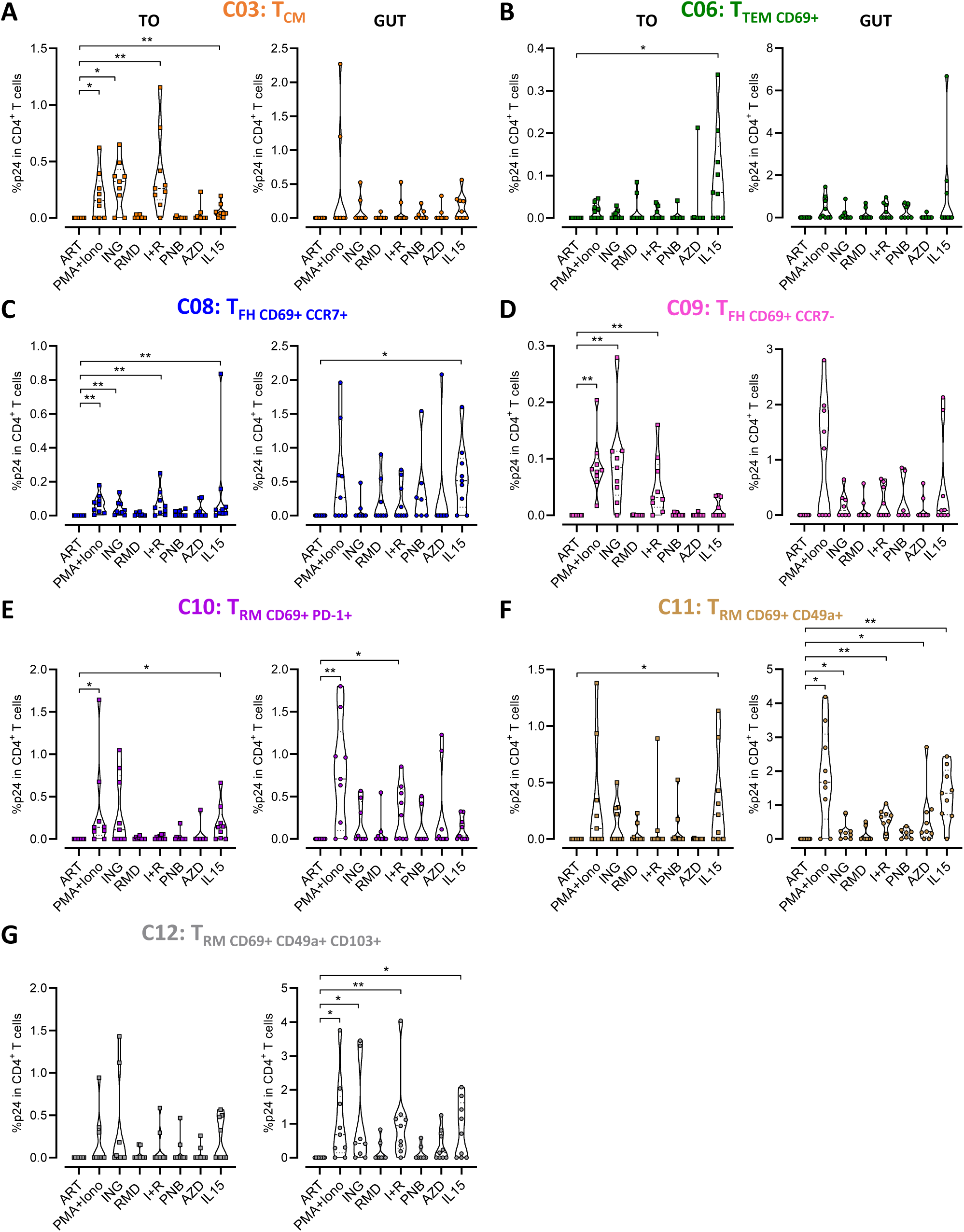
Heterogeneous responses of tonsillar and intestinal CD4^+^ T cell subpopulations to LRAs. Viral reactivation with LRAs in the ART-treated CD4^+^ T clusters from tonsils and intestines. Graphs displaying p24 expression within the CD4^+^ T cell subpopulations following stimulation with different LRAs: (A) C03:T_CM_, (B) C06:T_TEM CD69+_, (C) C08:T_FH CD69+ CCR7+_, (D) C09:T_FH CD69+ CCR7-_, (E) C10:T_RM CD69+ PD-1+_, (F) C11:T_RM CD69+ CD49a+_, and (G) C12:T_RM CD69+ CD49a+ CD103+_. Median values with quartiles are represented. Statistical comparisons of p24 levels between unstimulated (ART) and LRA-reactivated conditions were conducted using the Wilcoxon test, with significance levels denoted as *p < 0.05; **p < 0.01.

In summary, our findings reveal distinct reactivation patterns to current LRAs within the same CD4^+^ T cell subpopulations across the tissues. HIV reservoirs within resident T_FH_ subsets and T_CM_ cells exhibited superior responsiveness to current LRAs in the tonsils, whereas HIV within T_RM_ clusters showed greater reactivation in the intestine. This underscores the tissue-specific nature of both HIV reservoir establishment and reactivation dynamics.

## Discussion

Despite significant advancements in antiretroviral therapy, which have transformed HIV into a manageable chronic condition, a cure remains elusive. The persistence of HIV, mainly in long-lived CD4^+^ T cells where it can evade immune surveillance, poses a major barrier to eradication efforts^8^. These HIV reservoirs span diverse CD4^+^ T cell populations and anatomical compartments^19^, making it essential to understand their distinct characteristics and distribution to develop effective therapeutic strategies. In this study, we developed explant models of human tonsils, intestinal mucosa and cervicovaginal tissue to investigate the susceptibility of CD4^+^ T subpopulations within these tissues to HIV infection and potential reservoir establishment. We revealed tissue and cell-specific dynamics influencing HIV infection, viral persistence under ART, and HIV reactivation, demonstrating differential responses of the CD4^+^ T viral reservoirs to various LRAs. Our findings highlight the need for broad-spectrum therapies capable of effectively targeting the diverse cellular reservoirs of HIV across different anatomical compartments.

We developed explant models of HIV latency, based on an established experimental setup^88,89^, using three crucial tissues: tonsils, intestinal mucosa and cervix. The models accurately recapitulated key aspects of HIV infection. First, we observed a decline in CD4^+^ T cells during untreated HIV infection^90–92^. Second, ART administration partially reversed CD4^+^ T cell loss and facilitated viral reservoir establishment^5,93–95^. Third, ART caused a more pronounced decrease in intact HIV DNA compared to defective forms^96,97^. Fourth, we detected residual viral production despite ART, a phenomenon commonly observed in blood, lymph nodes and gastrointestinal tissue from PWH^98–102^. Although the first three observations could not be conclusively established in the cervix, residual viral transcription was detected in cervical mucosa despite ART suppression, consistent with prior reports from our group^9^. Overall, our explant models effectively mirror important events of both untreated and treated HIV infection in these tissues, providing valuable tools for investigating the mechanisms of HIV persistence and strategies directed to target tissue reservoirs.

In our study, we reported a predominance of T_NA_ and T_FH_ cells in tonsils and T_RM_ subsets in the intestinal mucosa and cervix, aligning with previous reports^9,47,54,73–75,78,79^. It is important to consider that age differences among tissue donors may influence these proportions and, as a result, impact the susceptibility to HIV infection of the tissues. Tonsils derived from paediatric individuals exhibit a higher proportion of less differentiated cells compared to those from adults^103^. Furthermore, aging is known to increase the frequency of T_RM_ cells in cervical tissue^79^. Similarly, intestinal samples from adult donors may display a higher prevalence of more differentiated and senescent cells due to the advanced age of the individuals.

Productive HIV infection was detected in the majority of the clusters in the tonsils and intestine but, remarkably, T_RM_ cells consistently exhibited the highest levels of HIV infection and significantly contributed to the pool of infected CD4^+^ T cells in both tissues. This was particularly noteworthy in the tonsils, where, despite being scarce (approximately 5% of total CD4^+^ T cells), T_RM_-phenotype infected cells were notably abundant comprising approximately 27% of total infected cells. Moreover, we have previously demonstrated a preferential infection of T_RM_ cells in cervical tissue^9^. Consequently, our findings, supported by other studies^104,105^, strongly suggest that T_RM_ cells are preferential targets for HIV infection within tissues. Furthermore, HIV infection has been reported to induce a T_RM_-like phenotype in resting memory CD4^+^ T cells^106^ which enhances their susceptibility to productive infection. This further supports the notion that the T_RM_-phenotype might be associated with increased permissiveness to HIV infection.

After ART administration, inducible viral reservoirs were identified in all tissue models. In tonsils, we detected HIV reservoirs across a broad range of cell subpopulations. Notably, resident T_FH_ cell clusters and the T_RM_ subset C10:T_RM CD69+ PD-1+_ were prominent contributors to the inducible HIV reservoir. This aligns with previous studies specifically linking CXCR5, PD-1 and CD69 with latent HIV infection in tonsillar tissue^15,107^. Furthermore, CD4^+^ T_CM_ cells, recognized as major HIV reservoirs in peripheral blood^21,59,61^, also play a significant role as inducible viral reservoirs within the tonsillar lymphoid tissue. In the intestine, T_RM_ cells, particularly those with high CD127 and slight KLRG1 expression, emerged as the predominant HIV reservoir. CD4^+^ T cells expressing CD69 and PD-1 have been previously shown to support latent HIV infection in the intestinal tissue^15^. Moreover, T_RM_ markers CD49a and CD127, as well as KLRG1 have been associated with CD4^+^ T cells harbouring HIV reservoirs^9,14,46,71^. Notably, our study identifies CD69, CD45RO and PD-1 as common markers of inducible reservoirs in both tissues, while CXCR5 is specific to tonsils and CD49a and CD103 are exclusive markers of HIV reservoirs in the intestine. These findings reveal a spectrum of phenotypes contributing to HIV persistence in tissues, with populations expressing markers previously associated with HIV latency, underscoring their potential as therapeutic targets.

Our study evaluated several LRAs for their capacity to reactivate HIV within tissue reservoirs. Such investigations with human tissues are infrequent, primarily due to the challenges involved in obtaining samples. Among the LRAs tested, Ingenol demonstrated great potency particularly in tonsillar and cervical reservoirs and broad reactivation activity across the distinct tonsillar CD4⁺ T cell subpopulations. In tonsils, Ingenol preferentially reactivated the T_CM_ subset over the other non-resident populations, mirroring similar patterns observed in peripheral blood^61^. Moreover, Ingenol showed robust reactivation of T_FH_ subsets in the tonsils and T_RM_ cells in the intestine. The differential responsiveness among these subsets may be attributed to varying levels of PKC isoforms and NF-κB pathway components. In contrast, both HDACis Romidepsin and Panobinostat, known for their ability to reactivate latent HIV in blood cells^60,61,63,108–110^, showed no effect on any tissue. Notably, Panobinostat’s ineffectiveness in reactivating tissue reservoirs has also been corroborated in animal models^108^. Previous research has demonstrated that these drugs differ in their ability to inhibit cell-associated HDAC activity^60^. Therefore, it is plausible to suggest that the differential expression of HDAC isoforms between the CD4^+^ T cell subsets in blood and tissues might contribute to their intrinsic limitations in reactivating HIV in tissues.

Recently, the potential use of IL15 as LRA has been extensively evaluated. IL15 has demonstrated efficacy in reactivating HIV^62,111–114^ and, unlike most LRAs, is capable of priming latently infected cells for recognition by CD8^+^ T cells and NK cells while also enhancing the cytotoxic activity of these cells^62,112,115^. Our study revealed a potent effect of IL15 in inducing HIV reactivation across all studied tissues, demonstrating efficacy even in CD4^+^ T cell subsets unresponsive to PMA/Ionomycin stimulation. The dual role of IL15 in both reactivating HIV and promoting cell survival and proliferation^116,117^ may account for the observed increase in p24 production. The heightened responsiveness of certain CD4⁺ T subsets to IL15 is likely attributable to increased expression of the IL-15 receptor (IL-15R). IL-15 dependence for CD8⁺ T_RM_ cell maintenance varies across tissues^118^, suggesting tissue-specific differences in IL-15R expression within these populations, a pattern that may extend to the CD4⁺ T_RM_ compartment. Although T_CM_ cells are generally not associated with elevated IL-15R expression^119^, our findings revealed IL-15-mediated reactivation in the tonsillar T_CM_ subset. Additionally, while T_FH_ cells from LNs have been reported to express higher levels of IL-15R compared to other memory subsets^46^, only select T_FH_ subpopulations responded to IL-15 stimulation. These observations suggest heterogeneity in IL-15R expression across tissues and within distinct CD4⁺ T cell subpopulations. Beyond receptor expression, IL15-induced intracellular signalling pathways may further modulate HIV reactivation. IL15 has been reported to inhibit the action of SAM domain and HD domain-containing protein 1 (SAMHD1)^117^, an enzyme that impairs HIV gene expression and negatively modulate viral latency reactivation in CD4^+^ T cells^120^. Notably, T_FH_ cells exhibit low level of SAMHD1^121^, which may facilitate their susceptibility to IL15-mediated reactivation. Collectively, these findings suggest that the differential responsiveness of CD4⁺ T cell subsets to IL15 may arise from the combined effects of IL15 receptor expression and SAMHD1 activity, a hypothesis that warrants further investigation.

AZD5582 has demonstrated significant efficacy in reactivating HIV in humanized mice and SIV-infected rhesus macaques under ART^122,123^. In these animal models, HIV reactivation was observed in lymph nodes^122^, but its impact in the gastrointestinal tract remained unexplored. Conversely, we revealed that AZD5582 exclusively induced HIV reactivation in the intestine, with minimal effect on CD4^+^ T cells from the tonsillar and cervical tissues. The differences in findings may arise from the detection techniques employed; while that study measured viral RNA, our detection relied on p24 protein levels which may selectively capture only the most robust reactivations^124,125^. Indeed, AZD5582’s effect was only observed in the T_RM_ subset with the highest levels of infection and susceptibility to viral reactivation by most LRAs studied. These observations suggest that AZD5582 may exhibit only a modest potency in CD4^+^ T cells form intestine, triggering viral reactivation primarily in the subsets harbouring more inducible HIV.

We observed that certain CD4^+^ T cell subsets in tonsillar and intestinal tissues, despite exhibiting high infection rates, showed resistance to viral reactivation. One possible explanation is that LRAs might enhance HIV RNA transcription in specific cells but fail to induce robust production of viral particles, as previously demonstrated in blood^124,125^. Another possibility is that these cell types tend to sustain productive rather than latent HIV infection and may undergo rapid cell death following infection^126,127^. Additionally, LRAs might induce phenotypic changes that hinder the identification of the original cell subset. Ultimately, specific phenotypes of tonsillar and intestinal CD4^+^ T cells could be in a “deeper” latent state than those in peripheral blood, rendering them less responsive to LRA stimulation. Indeed, this could be the case for intestinal CD4^+^ T cells, as elevated HIV transcription has been observed in circulating CD4^+^ T cells compared to rectal tissue CD4^+^ T cells following LRA stimulation. A greater block to HIV transcription has been described as responsible for this phenomenon^128,129^. Moreover, silencing due to repressive chromatin modifications or transcriptional interference^130–132^, as well as integration into heterochromatin^133^, could profoundly impact viral reactivation in specific cell subsets. Importantly, these mechanisms are largely unexplored in tissue cells. Overall, the absence of detectable viral reactivation in certain CD4^+^ T cell subsets susceptible to productive HIV infection may be attributed to a combination of factors, including defective production of viral particles, LRA-induced phenotypic changes, and profound transcriptional silencing within the cells.

Our study has several limitations. Firstly, the lifespan of the tissues is limited, preventing long-term studies of HIV persistence. However, since the majority of HIV reservoirs are established shortly after initial infection and at the onset of ART^5^, we consider our models well-suited for identifying potential cell reservoirs. Secondly, we used a CCR5-tropic HIV strain and did not test CXCR4-tropic viruses. While differences in infection rates and affected CD4^+^ T cell subpopulations may occur, we believe the inducibility of the established reservoirs would not be significantly altered. Furthermore, using a CCR5- tropic strain more accurately models an early HIV infection, making it more suitable for studying viral reservoir establishment. Thirdly, due to the difficulty in isolating individual tissue subsets for reservoir identification, we identified inducible reservoirs after maximal *in vitro* activation. While most non-inducible proviruses in blood of PWH on ART are likely defective, studies suggest that 11.7% may comprise intact HIV genomes^134^. Whether this fraction differs in tissue cells remains unclear. Additionally, it is uncertain if the inducible reservoirs identified in our study contain replication-competent provirus. Moreover, it remains unknown whether the observed increase in p24 expression detected by flow cytometry reflects truly viral reactivation from latently infected cells or enhanced viral protein production in previously undetectable virus-producing cells. Finally, a principal limitation of our study is the uncertainty regarding potential alterations in the CD4^+^ T cell phenotype following pharmacological reactivation^15,22,83–85^. However, we found that the frequency of the main clusters in both unstimulated and stimulated conditions remained consistent, as also observed in a previous study^59^. Moreover, we did not observe the creation of new cell clusters after viral reactivation as reported in an early study^15^.

In summary, using highly physiological tissue models, we identified inducible cellular HIV reservoirs in critical tissues with varying susceptibility to current LRAs. The differential responses to pharmacological viral reactivation among different CD4^+^ T cell subpopulations within tonsillar and intestinal tissues, as well as variations within the same subpopulations across the two tissues, suggest that viral reactivation is influenced by both cellular and tissue-specific factors. Therefore, our findings have significant implications for designing therapies aimed at promoting HIV reactivation in tissues.

## Materials and methods

### Patients samples and ethics statement

Tonsils were collected from routine tonsillectomies and tonsil reduction surgeries performed at the Otorhinolaryngology Department of Hospital Universitari Vall d’Hebron (HUVH). Gastrointestinal tissues, specifically colon resections, were procured from colorectal cancer patients undergoing colectomies at the Gastroenterology department of HUVH. Cervicovaginal tissues were obtained from non-neoplastic hysterectomies and cervical amputations performed in the Gynaecology Department of HUVH. The study protocols received ethical approval from the Institutional Review Board at HUVH [PR(AG)582/2020PR]. All tissues were surgically removed for non-inflammatory reasons, and only healthy regions of the collected samples were used in the study. Participants included paediatric patients for tonsillar samples and adult patients for intestinal and cervical tissues, all confirmed to be HIV-negative at the time of collection. Participation in the study required written informed consent provided by adult participants themselves or by legal guardians for paediatric patients. The anonymity and untraceability of the collected samples were ensured. Information on the number of participants, their gender and age is provided in **Table S1**.

### Tissue explant models of productive and latent HIV infection

Tonsillar, intestinal and cervicovaginal tissues were processed as previously described ^88,89^. Briefly, tissue resections were delivered in RPMI 1640 medium (Life Technologies, cat.no. 12004997) containing antibiotics and antifungals within 4 hours post-surgery. The tissues were dissected into uniform blocks of approximately 2 mm x 2 mm x 2 mm in size. For tonsillar and intestinal tissues, 10-12 blocks were placed on a piece of absorbable gelatine sponge (Surgispon, Aegis Lifesciences, cat. no. SSP-805010) suspended in RPMI 1640 medium supplemented with 20% fetal bovine serum (FBS, Life Technologies, cat. no. A5256701) (R20 medium), 100 U/ml penicillin, 100 µg/ml streptomycin (Capricorn Scientific, cat. no. PS-B), and 66 µg/ml Amikacin (Normon, cat. no. 791301.6) in a 6-well plate. Timentin (Caisson Labs, cat. no. T034) at a concentration of 310 µg/ml was added to the media only on the first day of culture. Viral infection was initiated by depositing 5 µl of the CCR5-tropic HIV viral strain HIV_BAL_ (156,250 TCID_50_) onto each tissue block, with some blocks left uninfected to serve as controls. Infected and control blocks were cultured for an additional 7-9 days at 37 °C and 5% CO_2_, with the media and sponges replaced every 3 days. On day 5 (tonsils) or day 6 (intestine), ART comprising 1 µM Raltegravir (NIH AIDS Reagent Program, cat. no. 0980), 1 µM Darunavir (NIH AIDS Reagent Program, cat. no. 0989), and 1 µM Nevirapine (Sigma-Aldrich, cat.no. SML0097), was added on top of the infected tissue blocks and left for a period of 2-3 days. For cervicovaginal explants, 8-12 blocks of tissue per condition were immersed in 310 µl R20 media with 100 U/ml penicillin and 100 µg/ml streptomycin in a 24-well plate and infected with 40 µl of HIV_BAL_ virus (156,250 TCID50). Some blocks were placed in 350 µl of medium to serve as controls. After a 2-hour incubation at 37 °C and 5% CO_2_, tissue blocks were rinsed three times with 3 mL of 1X PBS in 6 well plates and transferred back into a 12-well plate, placing 8-12 blocks per well in 1 mL of R20 media with antibiotics. Infected and control blocks were cultured for an additional 8 days, with the medium replaced every 3 days. On day 6, ART comprising 1 µM Raltegravir, 1 µM Darunavir, 1 µM and Nevirapine was added to the culture medium containing some of the infected tissue blocks for 2 days.

### Digestion of tissue blocks

Digestion procedures were optimized for each tissue based on established protocols ^88,89^. Tonsillar blocks were mechanically digested using disposable pellet pestles in RPMI 1640 supplemented with 5% FBS (R5). For intestinal and cervicovaginal tissues, enzymatic digestion was required prior to the mechanical disruption of the blocks. Intestinal blocks were incubated in R5 containing 2.5 mg/ml collagenase IV (Fisher Scientific, cat. no. 10780004) and 100 μg/ml DNase I (Roche, cat. no.10104159001) while cervicovaginal blocks were submerged in R5 with 5 mg/ml collagenase IV, both for 30 min at 37 °C and 400 rpm. Once digested, all cellular suspensions were filtered through 70 μm cell strainers (Labclinics, cat.no. PLC93070) to remove large aggregates. Finally, cells were washed twice with 1X PBS.

### Longitudinal phenotyping of key immune populations and HIV infection assessment by flow cytometry

To characterize changes in pivotal immune populations and measure HIV infection in CD4^+^ T cells during the culture of the infected explants, tissue blocks were digested at various time points and subsequently stained for both surface and intracellular markers. Cellular suspensions were stained with LIVE/DEAD Fixable Aqua Dead Cell Stain (Invitrogen, cat. no. L34966) for 20 min at room temperature (RT). After washing with 1X PBS, cells were incubated with anti-HLA-DR-PerCP-Cy5.5 (G46-6, Beckton Dickinson, cat. no. 560652), anti-CD56-FITC (B159; Beckton Dickinson, cat. no. 562794), anti-CD3- PE-Cy5 (UCHT-1, Biolegend, cat. no. 300410), anti-CD45-AF700 (HI30, Biolegend, cat. no. 304024), anti-CD8-APC (RPA-T8, Beckton Dickinson, cat. no. 555369), anti-CD16- BV605 (3G8, Beckton Dickinson, cat. no. 563172) and anti-CD19-V500 (HIB19, Beckton Dickinson, cat.no. 561121) antibodies in staining buffer (1X PBS 3% FBS) for 20 min at RT. After washing with staining buffer, cells were fixed and permeabilized using Fixation/Permeabilization Solution (Beckton Dickinson; cat. no. 554714;) for 20 min at 4°C, followed by two washes with BD Perm/Wash™ buffer. Then, intracellular staining with anti-p24 antibody (KC51, Beckman Coulter, cat. no. 6604667) was performed for 30 min on ice, followed by an additional 30 min at RT. After washing, cells were fixed with 2% PFA. Samples were acquired in a LSRFortessa (Becton Dickinson) flow cytometer and analysed with FlowJo v10 software (TreeStar).

The degree of HIV infection in CD4^+^ T cells was also assessed by staining digested tissue blocks using a panel comprising: LIVE/DEAD™ Fixable Far Red Dead Cell Stain (Invitrogen, cat. no. L34974), anti-CD45-FITC (HI30, Biolegend, cat. no. 304006) or anti-CD45-BV605 (HI30, Becton Dickinson, cat. no. 564047), anti-CD3-Per-Cp (SK7, Becton Dickinson, cat. no. 345766) or anti-CD3-AF700 (SK7, Biolegend, cat. no. 344822), anti-CD8-APC (RPA-T8, Beckton Dickinson, cat. no. 555369) and anti-p24-PE (KC51, Beckman Coulter, cat. no. 6604667) antibodies. Viability, surface, and intracellular staining, as well as fixation, were performed as previously described. Sample acquisition was conducted using a BD FACSCalibur or a BD LSRFortessa flow cytometer, and data analysis was carried out with FlowJo v10 software.

### Quantification of HIV DNA in isolated CD4^+^ T cells by Intact proviral DNA assay (IPDA) and Quantitative PCR (qPCR)

To quantify all forms of HIV DNA within infected explants, tissue blocks from tonsils collected on day 7, and from intestinal and cervicovaginal tissues collected on day 8, were digested as previously described. Following digestion, cellular suspensions were processed with the EasySep™ Dead Cell Removal (Annexin V) Kit (StemCell, cat. no. 17899) to remove apoptotic cells. Two rounds of immunomagnetic negative selection were performed to ensure isolation of only live cells. Subsequently, CD4^+^ T cells were isolated from the live cell fraction using the Dynabeads™ CD4 Positive Isolation Kit (Invitrogen, cat. no. 11331D). Isolated cells were lysed in a Proteinase K-containing lysis buffer at 55°C over-night and at 95°C for 5 minutes. IPDA was then conducted on lysed CD4^+^ T cells from tonsillar and intestinal tissues according to the established protocol^135^, using specific primers and probes targeting the Ψ HIV gene (HIV-1 Ψ forward 50 5’-CAGGACTCGGCTTGCTGAAG-3’, HIV-1 Ψ reverse 5’-GCACCCATCTCTCTCCTTCTAGC-3’ and probe 5’ 6-FAM-TTTTGGCGTACTCACCAGT-MGBNFQ-3’) and the env HIV gene (HIV-1 env forward 5’-AGTGGTGCAGAGAGAAAAAAGAGC-3’, HIV-1 env reverse 5’-GTCTGGCCTGTACCGTCAGC-3’, HIV-1 env intact probe 5’-VIC-CCTTGGGTTCTTGGGA-MGBNFQ-3’, and HIV-1 anti-Hypermutant *env* probe 5’- CCTTAGGTTCTTAGGAGC-MGBNFQ-3’). The hRPP30 gene served as a reference for cell input normalization. Samples were analysed in a QIAcuity One 2 plex System (Qiagen). IPDA could not be performed on isolated CD4^+^ T cells from cervicovaginal tissue due to the limited cell yield. Consequently, qPCR was employed as an alternative method. Total HIV DNA in the CD4^+^ T cell lysates was quantified by qPCR using primers and probes specific for the 1-LTR HIV region (LTR forward: 5’- TTAAGCCTCAATAAAGCTTGCC-3’, LTR reverse: 5’-GTTCGGGCGCCACTGCTAG-3’, and probe: 5’ /56-FAM/CCAGAGTCA/ZEN/CACAACAGACGGGCA/ 31ABkFQ/ 3’). The CCR5 gene was used for cell input normalization. Samples were analysed in a QuantStudio^TM^ 5 Real-Time PCR System, and quantification of DNA copies was performed using a standard curve.

### E*x vivo* reactivation with LRAs of HIV-infected and ART-treated CD4^+^ T cells

Tissue blocks from uninfected, HIV-infected, and HIV-infected and ART-treated conditions collected on day 7 for tonsils, and on day 8 for the intestinal and cervicovaginal tissue, were digested according to the previously described protocol. CD4^+^ T cells were then isolated using the MagniSort Human CD4^+^ T Cell Enrichment commercial kit (Invitrogen, cat. no. 8804-6811-74), with two rounds of cell separation performed to maximize cell purity. In all conditions, cells were cultured in R10 media containing 10 μM Q-VD-OPh (Selleckchem, cat. no. S7311) for a minimum of 2 h at 37°C and 5% CO_2_. For HIV-infected and ART-treated CD4^+^ T cells, 1 µM Raltegravir, 1 µM Darunavir and 1 µM Nevirapine were added to the R10 media. After the incubation, uninfected and HIV-infected CD4^+^ T cells remained untreated, while HIV-infected and ART-treated CD4^+^ T cells were stimulated for 20-22 hours with latency reversal agents (LRAs). LRAs were added to the media at the following concentrations: 100 nM Ingenol-3-angelate (Sigma-Aldrich, cat. no. SML1318), 40 nM Romidepsin (Selleckchem, cat. no. S3020), 30 nM Panobinostat (Selleckchem, cat. no. S1030), 100 nM AZD5582 (MedChem, cat.no. HY- 12600), 50 ng ml^-1^ IL15 (Myltenyi Biotech, cat.no. 130-095-760). The positive control consisted of 81 nM PMA and 1 μM Ionomycin (Abcam, cat. no. ab120297 and ab120370) in the medium, while the negative control comprised solely of R10 medium with ART. All compounds were reconstituted in either deionized sterile-filtered water, 0.1% BSA in 1X PBS, or DMSO (at a maximum concentration of 0.006%).

### CD4^+^ T subsets phenotyping and detection of p24 protein by flow cytometry

Identification of CD4^+^ T subpopulations present in the tissues and measurement of HIV infection through p24 protein detection within these subsets was performed using flow cytometry. Previously cultured CD4^+^ T cells were stained using LIVE/DEAD Fixable Aqua Dead Cell Stain (Invitrogen, cat. no. L34966) for 20 min at RT. Following washing with 1X PBS, cells were stained with anti-CXCR5-BB700 (RF8B2, Beckton Dickinson, cat. no. 566470) and anti-CCR7-AF647(150503, Beckton Dickinson, cat. no. 560816) in staining buffer for 30 min at 37°C. After washing with staining buffer, cells were incubated with the following antibodies for 20 min at RT: anti-CD127-FITC (A019D5, Biolegend, cat. no. 351312), anti-CD49a-PE-Cy7 (TS2/7, Biolegend, cat. no. 328311), anti-CD3-PE- Cy5 (UCHT-1, Biolegend, cat. no. 300410) anti-CD69-PE-CF594 (FN50, Beckton Dickinson, cat. no. 562617), anti-PD-1-APC-Fire750 (EH12.2H7, Biolegend, cat. no. 329954), anti-CD45-AF700 (HI30, Biolegend, cat. no. 304024), anti-CD103-BV650 (Ber-ACT8, Beckton Dickinson, cat. no. 743653), anti-CD45RO-BV605 (UCHL1, Biolegend, cat. no. 304238) and anti-KLRG1-BV421 (14C2A07, Biolegend, cat. no. 368604). Next, cells were washed with staining buffer and fixed and permeabilized using Fixation/Permeabilization Solution (Beckton Dickinson; cat. no. 554714) for 20 min at 4°C. Two washes with BD Perm/Wash™ buffer were conducted and intracellular staining with anti-p24 antibody (KC51, Beckman Coulter, cat. no. 6604667) was then performed for 30 min on ice, followed by an additional 30 min at RT. Cells were then washed and fixed with 2% PFA. Samples were acquired using a BD LSR Fortessa flow cytometer.

The acquired data were analysed using the online platform OMIQ (www.omiq.ai, www.dotmatics.com). The clustering algorithm Flow-Self Organizing Maps (FlowSOM) was employed to identify clusters based on the expression of CD45RO, CCR7, CD69, CD103, CD49a, CD127, CXCR5, PD-1 and KLRG1. All events within pre-gated live CD3^+^ cells were concatenated for each group and subsequently analyzed. To visualize the identified clusters, the dimensionality reduction algorithm opt-SNE was applied. Opt-SNE was performed on a randomly selected subset of 6 x 10^5^ cells from uninfected tonsillar and intestinal tissues and 4 x 10^3^ cells (all cells available) from cervical tissue based on the expression of the aforementioned markers. Additionally, FlowJo v10 software was used to apply Boolean filters to identify, based on the expression of mentioned markers, the specific subsets of live CD3^+^ p24^+^ cells in the PMA/Ionomycin-stimulated conditions from tonsillar and intestinal tissues.

### Statistical analyses

All data were analysed using GraphPad Prism 8.3.0 software (GraphPad Software, La Jolla, CA, USA), with statistical significance defined as p< 0.05. Volcano plots, generated using OMIQ software (www.omiq.ai, www.dotmatics.com), represent statistical comparisons conducted via the edgeR method, with significance determined at p< 0.05. Detailed statistical information for each experiment is provided in the respective figure legends.

## Author contributions

AGC and MJB designed, directed, and interpreted experiments. AGC, NSG, CMP, ORI, and ABM performed the experiments and analysed the data. MG contributed to the design and interpretation of the experiments. JGP, ML and RP produced and provided the HIV_BAL_ viral stock. SL, JC, FP, NO, IL and JL were responsible for recruitment, specimen handling and storage, and the collection of related clinical data. AGC and MJB wrote the initial manuscript, and all authors contributed to its editing.

## Acknowledgments

This study was funded by “la Caixa” Foundation (ID 100010434) under the project code LCF/PR/HR20/00218. This study also received funding from the Agencia Estatal de Investigación project PID2021-123321OB-I00 supported by MCIN/AEI/10.13039/501100011033/FEDER, UE; The Spanish “Ministerio de Economía y Competitividad, Instituto de Salud Carlos III” (ISCIII, PI20/00160); and the Gilead fellowships GLD21-00049 and GLD22/00152. MJB is supported by the Miguel Servet program of the Spanish Health Institute Carlos III (CP17/00179 and CPII22/00005), and AGC is a recipient of a PhD fellowship from “la Caixa” Foundation (LCF/BQ/DI20/11780014). The funders had no role in the design of the study, data collection or analysis, decision to publish, or manuscript preparation.

## Supplementary Figure legends

**Supplementary Figure 1. Identification of key immune populations in the tonsillar and intestinal models of latent HIV infection.** Related to Figure 1.

(A) Representative example of the gating strategy used for the identification of CD4^+^ T cells, CD8^+^ T cells, NK cell populations (total CD56^+^, CD56^+^ CD16^+^ and CD56^+^ CD16^-^ NK cells) and B cells; as well as the assessment of HIV infection levels based on p24 expression within CD4^+^ T cells in tonsillar and intestinal tissues.

(B) Fluorescence Minus One (FMO) controls for markers used in the identification of B cells and NK cells. FMOs for markers HLA-DR and CD19, CD56 and CD16 are gated within the live CD45^+^ cells. FMOs for markers CD56 and CD16 are gated within CD45^+^ CD3^-^ CD8^-^ live cells.

**Supplementary Figure 2. Longitudinal monitoring of key immune populations during productive and treated HIV infection.** Graphs illustrating the frequency of CD4^+^ T cells, CD8^+^ T cells, total CD56^+^ NK cells, CD56^+^ CD16^+^ NK cells, CD56 ^+^ CD16^-^ NK cells and B cells within CD45^+^ live cells in the uninfected, infected and infected and ART-treated conditions in both tonsillar and intestinal tissues over an 8-9-day period. Means and standard error of means (SEM) are depicted. Related to Figure 1.

**Supplementary Figure 3. Expression of the markers used in the identification of the CD4^+^ T cell clusters.** Related to Figure 2.

(A) Histograms displaying the Mean Fluorescence Intensity (MFI) for the markers used to identify the twelve CD4^+^ T cell clusters. Each cluster is represented in a different color, corresponding to Figure 2.

(B) Fluorescence Minus One (FMO) controls for the markers CD45RO, CCR7, CD69, CD103, CD49a, CXCR5, PD-1, CD127, and KLRG1. FMO controls are gated within live CD45^+^ CD3^+^ cells, showing background fluorescence and allowing for accurate gating of the CD4^+^ T cell clusters.

**Supplementary Figure 4. Productive HIV infection in CD4^+^ T clusters of intestinal and tonsillar tissues.** Related to Figure 2.

(A) Volcano plot illustrating differences in the proportion of CD4^+^ T cell clusters between infected conditions in tonsillar and intestinal tissues. Statistically significant differences are highlighted by green dots.

(B) Tonsillar and intestinal CD4^+^ T cell compartments in the HIV-infected condition. Pies charts depicting the medians of the percentages of each cluster within the total CD4^+^ T cells from infected tonsil and intestine. Statistical values derived from the Volcano plot in (A) with significance levels indicated as *p < 0.05; **p < 0.01 and ****p < 0.0001

(C) Differences in cluster contributions to overall HIV infection levels. Graphs show the percentage contribution of each cluster to the total population of infected CD4^+^ T cells in both tissues. Statistical comparisons were performed using the Friedman test.

Statistical significance levels are represented as *p < 0.05; **p < 0.01; ***p < 0.001 and ****p < 0.0001.

**Supplementary Figure 5. Identification of inducible HIV reservoirs in the infected and ART-treated CD4^+^ T subpopulations from tonsillar and intestinal tissues.** Related to Figure 3.

(A) Volcano Plots and pie charts illustrating differences in cluster proportions between the ART-treated (ART) and ART-treated PMA-Ionomycin stimulated conditions in the intestine. Green dots in the Volcano plot indicate statistically significant differences. Pies charts depict the medians of the percentages of each cluster within the total CD4^+^ T cells. Statistical values derived from the Volcano plot.

(B) Volcano Plots and pie charts illustrating differences in cluster proportions between the ART-treated and ART-treated PMA-Ionomycin stimulated conditions in the tonsil. Green dots in the Volcano plot indicate statistically significant differences. Pies charts depict the medians of the percentages of each cluster within the total CD4^+^ T cells. Statistical values derived from the Volcano plot.

(C) Contribution of each cluster to overall inducible reservoir levels in CD4^+^ T cells within each tissue. Percentages are depicted and statistical comparisons were conducted using the Friedman test.

Significance levels are indicated as *p < 0.05; **p < 0.01; ***p < 0.001 and ****p < 0.0001.

**Supplementary Figure 6. Differences in the efficacy of current LRAs in inducing viral reactivation of tonsillar and intestinal HIV reservoirs.** Related to Figure 4.

(A-B) Volcano plots showing the differences in the proportion of clusters between the ART-treated (ART) and ART-treated and LRA-reactivated conditions in tonsillar (A) and intestinal (B) tissues. Green dots indicate statistically significant differences. The edgeR statistical method is employed, with green dots indicating statistically significant differences.

(C) Effect of Romidepsin (RMD) and Panobinostat (PNB) on HIV reactivation in tonsillar and intestinal clusters. Graphs displaying p24 expression within infected and ART-treated CD4^+^ T cells following stimulation with RMD and PNB. Median values with quartiles are represented. Statistical comparisons were performed using the Wilcoxon test.

**Supplementary Figure 7. Heterogeneous responses of tonsillar and intestinal CD4^+^ T cell subpopulations to LRAs.** Related to Figure 5. Bar chart displaying the potency of reactivation with IL15 compared to PMA and ionomycin, represented as fold change in C08:T_FH CD69+ CCR7+_ and C11:T_RM CD69+ CD49a+_ from tonsillar and intestinal tissues. Mean values with Standard Error of the Mean (SEM) are depicted. Statistical comparisons were performed using the Mann Whitney test.

## Supplementary Table legend

**Supplementary Table 1. Donor characteristics of human tissue samples.** This table summarizes the donor information for the human tissue samples used in the study, including the number of donors per tissue type, sex and age.

## References

1. Palmer, S., Josefsson, L. & Coffin, J. M. HIV reservoirs and the possibility of a cure for HIV infection. J Intern Med 270, 550–560 (2011).

2. Coiras, M., López-Huertas, M. R., Pérez-Olmeda, M. & Alcamí, J. Understanding HIV-1 latency provides clues for the eradication of long-term reservoirs. Nat Rev Microbiol 7, 798–812 (2009).

3. Gantner, P. et al. HIV rapidly targets a diverse pool of CD4+ T cells to establish productive and latent infections. Immunity 56, 653–668.e5 (2023).

4. Tokarev, A. et al. Single-Cell Profiling of Latently SIV-Infected CD4 ^+^ T Cells Directly *Ex Vivo* to Reveal Host Factors Supporting Reservoir Persistence. Microbiol Spectr 10, (2022).

5. Abrahams, M.-R. et al. The replication-competent HIV-1 latent reservoir is primarily established near the time of therapy initiation. Sci Transl Med 11, (2019).

6. Finzi, D. et al. Identification of a Reservoir for HIV-1 in Patients on Highly Active Antiretroviral Therapy. Science (1979) 278, 1295–1300 (1997).

7. Lichterfeld, M., Gao, C. & Yu, X. G. An ordeal that does not heal: understanding barriers to a cure for HIV-1 infection. Trends Immunol 43, 608–616 (2022).

8. Castro-Gonzalez, S., Colomer-Lluch, M. & Serra-Moreno, R. Barriers for HIV Cure: The Latent Reservoir. AIDS Res Hum Retroviruses 34, 739–759 (2018).

9. Cantero-Pérez, J. et al. Resident memory T cells are a cellular reservoir for HIV in the cervical mucosa. Nat Commun 10, 4739 (2019).

10. Serra-Peinado, C. et al. Expression of CD20 after viral reactivation renders HIV-reservoir cells susceptible to Rituximab. Nat Commun 10, 3705 (2019).

11. Gálvez, C. et al. Atlas of the HIV-1 Reservoir in Peripheral CD4 T Cells of Individuals on Successful Antiretroviral Therapy. mBio 12, (2021).

12. Grau-Expósito, J. et al. A Novel Single-Cell FISH-Flow Assay Identifies Effector Memory CD4 ^+^ T cells as a Major Niche for HIV-1 Transcription in HIV-Infected Patients. mBio 8, (2017).

13. Fromentin, R. et al. CD4+ T Cells Expressing PD-1, TIGIT and LAG-3 Contribute to HIV Persistence during ART. PLoS Pathog 12, e1005761 (2016).

14. Hsiao, F. et al. Tissue memory CD4+ T cells expressing IL-7 receptor-alpha (CD127) preferentially support latent HIV-1 infection. PLoS Pathog 16, e1008450 (2020).

15. Neidleman, J. et al. Phenotypic analysis of the unstimulated in vivo HIV CD4 T cell reservoir. Elife 9, (2020).

16. Astorga-Gamaza, A. et al. Identification of HIV-reservoir cells with reduced susceptibility to antibody-dependent immune response. Elife 11, (2022).

17. Pieren, D. K. J., Benítez-Martínez, A. & Genescà, M. Targeting HIV persistence in the tissue. Curr Opin HIV AIDS 19, 69–78 (2024).

18. Kandathil, A. J., Sugawara, S. & Balagopal, A. Are T cells the only HIV-1 reservoir? Retrovirology 13, 86 (2016).

19. Chen, J. et al. The reservoir of latent HIV. Front Cell Infect Microbiol 12, (2022).

20. Bosque, A. & Planelles, V. Studies of HIV-1 latency in an ex vivo model that uses primary central memory T cells. Methods 53, 54–61 (2011).

21. Chomont, N. et al. HIV reservoir size and persistence are driven by T cell survival and homeostatic proliferation. Nat Med 15, 893–900 (2009).

22. Kulpa, D. A. et al. Differentiation into an Effector Memory Phenotype Potentiates HIV-1 Latency Reversal in CD4 ^+^ T Cells. J Virol 93, (2019).

23. Hiener, B. et al. Identification of Genetically Intact HIV-1 Proviruses in Specific CD4 + T Cells from Effectively Treated Participants. Cell Rep 21, 813–822 (2017).

24. De Scheerder, M.-A. et al. HIV Rebound Is Predominantly Fueled by Genetically Identical Viral Expansions from Diverse Reservoirs. Cell Host Microbe 26, 347–358.e7 (2019).

25. Buzon, M. J. et al. HIV-1 persistence in CD4+ T cells with stem cell–like properties. Nat Med 20, 139–142 (2014).

26. Venanzi Rullo, E., et al. Persistence of an intact HIV reservoir in phenotypically naive T cells. JCI Insight 5, (2020).

27. Chaillon, A. et al. HIV persists throughout deep tissues with repopulation from multiple anatomical sources. Journal of Clinical Investigation 130, 1699–1712 (2020).

28. Wong, J. K. & Yukl, S. A. Tissue reservoirs of HIV. Curr Opin HIV AIDS 11, 362– 370 (2016).

29. Yukl, S. A. et al. Differences in HIV Burden and Immune Activation within the Gut of HIV-Positive Patients Receiving Suppressive Antiretroviral Therapy. J Infect Dis 202, 1553–1561 (2010).

30. Jenabian, M.-A. et al. Immune tolerance properties of the testicular tissue as a viral sanctuary site in ART-treated HIV-infected adults. AIDS 30, 2777–2786 (2016).

31. Fukazawa, Y. et al. B cell follicle sanctuary permits persistent productive simian immunodeficiency virus infection in elite controllers. Nat Med 21, 132–139 (2015).

32. Persidsky, Y. & Poluektova, L. Immune privilege and HIV-1 persistence in the CNS. Immunol Rev 213, 180–194 (2006).

33. Fletcher, C. V. et al. Persistent HIV-1 replication is associated with lower antiretroviral drug concentrations in lymphatic tissues. Proceedings of the National Academy of Sciences 111, 2307–2312 (2014).

34. Rosen, E. P. et al. Antiretroviral drug exposure in lymph nodes is heterogeneous and drug dependent. J Int AIDS Soc 25, (2022).

35. Lamers, S. L. et al. HIV DNA Is Frequently Present within Pathologic Tissues Evaluated at Autopsy from Combined Antiretroviral Therapy-Treated Patients with Undetectable Viral Loads. J Virol 90, 8968–8983 (2016).

36. Dufour, C. et al. Near full-length HIV sequencing in multiple tissues collected postmortem reveals shared clonal expansions across distinct reservoirs during ART. Cell Rep 42, 113053 (2023).

37. McManus, W. R. et al. HIV-1 in lymph nodes is maintained by cellular proliferation during antiretroviral therapy. Journal of Clinical Investigation 129, 4629–4642 (2019).

38. Wu, V. H. et al. Assessment of HIV-1 integration in tissues and subsets across infection stages. JCI Insight 5, (2020).

39. Pantaleo, G. et al. Lymphoid organs function as major reservoirs for human immunodeficiency virus. Proceedings of the National Academy of Sciences 88, 9838–9842 (1991).

40. Pantaleo, G. et al. HIV infection is active and progressive in lymphoid tissue during the clinically latent stage of disease. Nature 362, 355–358 (1993).

41. Chun, T. et al. Persistence of HIV in Gut-Associated Lymphoid Tissue despite Long-Term Antiretroviral Therapy. J Infect Dis 197, 714–720 (2008).

42. Poles, M. A. et al. Lack of Decay of HIV-1 in Gut-Associated Lymphoid Tissue Reservoirs in Maximally Suppressed Individuals. JAIDS Journal of Acquired Immune Deficiency Syndromes 43, 65–68 (2006).

43. Perreau, M. et al. Follicular helper T cells serve as the major CD4 T cell compartment for HIV-1 infection, replication, and production. Journal of Experimental Medicine 210, 143–156 (2013).

44. Renault, C. et al. Th17 CD4+ T-Cell as a Preferential Target for HIV Reservoirs. Front Immunol 13, (2022).

45. Miles, B. & Connick, E. T FH in HIV Latency and as Sources of Replication-Competent Virus. Trends Microbiol 24, 338–344 (2016).

46. Sun, W. et al. Phenotypic signatures of immune selection in HIV-1 reservoir cells. Nature 614, 309–317 (2023).

47. Kumar, B. V. et al. Human Tissue-Resident Memory T Cells Are Defined by Core Transcriptional and Functional Signatures in Lymphoid and Mucosal Sites. Cell Rep 20, 2921–2934 (2017).

48. Szabo, P. A. et al. Single-cell transcriptomics of human T cells reveals tissue and activation signatures in health and disease. Nat Commun 10, 4706 (2019).

49. Poon, M. M. L. et al. Tissue adaptation and clonal segregation of human memory T cells in barrier sites. Nat Immunol 24, 309–319 (2023).

50. Shiow, L. R. et al. CD69 acts downstream of interferon-α/β to inhibit S1P1 and lymphocyte egress from lymphoid organs. Nature 440, 540–544 (2006).

51. Thome, J. J. C. et al. Spatial map of human T cell compartmentalization and maintenance over decades of life. Cell 159, 814–28 (2014).

52. Schenkel, J. M. & Masopust, D. Tissue-Resident Memory T Cells. Immunity 41, 886–897 (2014).

53. Park, S. L. et al. Local proliferation maintains a stable pool of tissue-resident memory T cells after antiviral recall responses. Nat Immunol 19, 183–191 (2018).

54. Wong, M. T. et al. A High-Dimensional Atlas of Human T Cell Diversity Reveals Tissue-Specific Trafficking and Cytokine Signatures. Immunity 45, 442–456 (2016).

55. Yokoi, T. et al. Identification of a unique subset of tissue-resident memory CD4 ^+^ T cells in Crohn’s disease. Proceedings of the National Academy of Sciences 120, (2023).

56. Reilly, E. C. et al. T _RM_ integrins CD103 and CD49a differentially support adherence and motility after resolution of influenza virus infection. Proceedings of the National Academy of Sciences 117, 12306–12314 (2020).

57. Kim, Y., Anderson, J. L. & Lewin, S. R. Getting the “Kill” into “Shock and Kill”: Strategies to Eliminate Latent HIV. Cell Host Microbe 23, 14–26 (2018).

58. Board, N. L., Moskovljevic, M., Wu, F., Siliciano, R. F. & Siliciano, J. D. Engaging innate immunity in HIV-1 cure strategies. Nat Rev Immunol 22, 499–512 (2022).

59. Pardons, M., Fromentin, R., Pagliuzza, A., Routy, J.-P. & Chomont, N. Latency-Reversing Agents Induce Differential Responses in Distinct Memory CD4 T Cell Subsets in Individuals on Antiretroviral Therapy. Cell Rep 29, 2783–2795.e5 (2019).

60. Wei, D. G. et al. Histone Deacetylase Inhibitor Romidepsin Induces HIV Expression in CD4 T Cells from Patients on Suppressive Antiretroviral Therapy at Concentrations Achieved by Clinical Dosing. PLoS Pathog 10, e1004071 (2014).

61. Grau-Expósito, J. et al. Latency reversal agents affect differently the latent reservoir present in distinct CD4+ T subpopulations. PLoS Pathog 15, e1007991 (2019).

62. Covino, D. A., Desimio, M. G. & Doria, M. Impact of IL-15 and latency reversing agent combinations in the reactivation and NK cell-mediated suppression of the HIV reservoir. Sci Rep 12, 18567 (2022).

63. Banga, R., Procopio, F. A., Cavassini, M. & Perreau, M. *In Vitro* Reactivation of Replication-Competent and Infectious HIV-1 by Histone Deacetylase Inhibitors. J Virol 90, 1858–1871 (2016).

64. Shi, J. et al. PD-1 Controls Follicular T Helper Cell Positioning and Function. Immunity 49, 264–274.e4 (2018).

65. Hardtke, S., Ohl, L. & Förster, R. Balanced expression of CXCR5 and CCR7 on follicular T helper cells determines their transient positioning to lymph node follicles and is essential for efficient B-cell help. Blood 106, 1924–1931 (2005).

66. Haynes, N. M. et al. Role of CXCR5 and CCR7 in Follicular Th Cell Positioning and Appearance of a Programmed Cell Death Gene-1High Germinal Center-Associated Subpopulation. The Journal of Immunology 179, 5099–5108 (2007).

67. Asrir, A., Aloulou, M., Gador, M., Pérals, C. & Fazilleau, N. Interconnected subsets of memory follicular helper T cells have different effector functions. Nat Commun 8, 847 (2017).

68. Huster, K. M. et al. Selective expression of IL-7 receptor on memory T cells identifies early CD40L-dependent generation of distinct CD8 ^+^ memory T cell subsets. Proceedings of the National Academy of Sciences 101, 5610–5615 (2004).

69. Robbins, S. H., Terrizzi, S. C., Sydora, B. C., Mikayama, T. & Brossay, L. Differential regulation of killer cell lectin-like receptor G1 expression on T cells. J Immunol 170, 5876–85 (2003).

70. Voehringer, D., Koschella, M. & Pircher, H. Lack of proliferative capacity of human effector and memory T cells expressing killer cell lectinlike receptor G1 (KLRG1). Blood 100, 3698–702 (2002).

71. Astorga-Gamaza, A. et al. KLRG1 expression on natural killer cells is associated with HIV persistence, and its targeting promotes the reduction of the viral reservoir. Cell Rep Med 4, 101202 (2023).

72. Nave, H., Gebert, A. & Pabst, R. Morphology and immunology of the human palatine tonsil. Anat Embryol (Berl) 204, 367–373 (2001).

73. Mueller, S. N., Gebhardt, T., Carbone, F. R. & Heath, W. R. Memory T Cell Subsets, Migration Patterns, and Tissue Residence. Annu Rev Immunol 31, 137– 161 (2013).

74. Hagel, J. P. et al. Defining T Cell Subsets in Human Tonsils Using ChipCytometry. The Journal of Immunology 206, 3073–3082 (2021).

75. Crotty, S. Follicular Helper CD4 T Cells (T _FH_). Annu Rev Immunol 29, 621–663 (2011).

76. Thornhill, J. P., Fidler, S., Klenerman, P., Frater, J. & Phetsouphanh, C. The Role of CD4+ T Follicular Helper Cells in HIV Infection: From the Germinal Center to the Periphery. Front Immunol 8, (2017).

77. Sathaliyawala, T. et al. Distribution and Compartmentalization of Human Circulating and Tissue-Resident Memory T Cell Subsets. Immunity 38, 187–197 (2013).

78. Woodward Davis, A. S., et al. The human memory T cell compartment changes across tissues of the female reproductive tract. Mucosal Immunol 14, 862 (2021).

79. Shen, Z., vom Steeg, L. G., Patel, M. V., Rodriguez-Garcia, M. & Wira, C. R. Impact of aging on the frequency, phenotype, and function of CD4+ T cells in the human female reproductive tract. Front Immunol 15, 1465124 (2024).

80. Xie, G. et al. Characterization of HIV-induced remodeling reveals differences in infection susceptibility of memory CD4+ T cell subsets in vivo. Cell Rep 35, 109038 (2021).

81. Matheson, N. J. et al. Cell Surface Proteomic Map of HIV Infection Reveals Antagonism of Amino Acid Metabolism by Vpu and Nef. Cell Host Microbe 18, 409–423 (2015).

82. Cavrois, M. et al. Mass Cytometric Analysis of HIV Entry, Replication, and Remodeling in Tissue CD4+ T Cells. Cell Rep 20, 984–998 (2017).

83. Cohn, L. B. et al. Clonal CD4+ T cells in the HIV-1 latent reservoir display a distinct gene profile upon reactivation. Nat Med 24, 604–609 (2018).

84. Pardons, M. et al. Single-cell characterization and quantification of translation-competent viral reservoirs in treated and untreated HIV infection. PLoS Pathog 15, e1007619 (2019).

85. Baxter, A. E. et al. Single-Cell Characterization of Viral Translation-Competent Reservoirs in HIV-Infected Individuals. Cell Host Microbe 20, 368–380 (2016).

86. Bui, J. K. et al. Ex vivo activation of CD4+ T-cells from donors on suppressive ART can lead to sustained production of infectious HIV-1 from a subset of infected cells. PLoS Pathog 13, e1006230 (2017).

87. Jubel, J. M., Barbati, Z. R., Burger, C., Wirtz, D. C. & Schildberg, F. A. The Role of PD-1 in Acute and Chronic Infection. Front Immunol 11, (2020).

88. Grivel, J.-C. & Margolis, L. Use of human tissue explants to study human infectious agents. Nat Protoc 4, 256–269 (2009).

89. Cantero, J. & Genescà, M. Maximizing the immunological output of the cervicovaginal explant model. J Immunol Methods 460, 26–35 (2018).

90. Grossman, Z., Meier-Schellersheim, M., Paul, W. E. & Picker, L. J. Pathogenesis of HIV infection: what the virus spares is as important as what it destroys. Nat Med 12, 289–295 (2006).

91. Veazey, R. S. et al. Gastrointestinal Tract as a Major Site of CD4 ^+^ T Cell Depletion and Viral Replication in SIV Infection. Science (1979) 280, 427–431 (1998).

92. Brenchley, J. M. et al. CD4+ T Cell Depletion during all Stages of HIV Disease Occurs Predominantly in the Gastrointestinal Tract. J Exp Med 200, 749–759 (2004).

93. Suanzes, P. et al. Impact of very early antiretroviral therapy during acute HIV infection on long-term immunovirological outcomes. International Journal of Infectious Diseases 136, 100–106 (2023).

94. Le, T. et al. Enhanced CD4+ T-Cell Recovery with Earlier HIV-1 Antiretroviral Therapy. New England Journal of Medicine 368, 218–230 (2013).

95. Schuetz, A. et al. Initiation of ART during Early Acute HIV Infection Preserves Mucosal Th17 Function and Reverses HIV-Related Immune Activation. PLoS Pathog 10, e1004543 (2014).

96. Gandhi, R. T. et al. Selective Decay of Intact HIV-1 Proviral DNA on Antiretroviral Therapy. J Infect Dis 223, 225–233 (2021).

97. Peluso, M. J., et al. Differential decay of intact and defective proviral DNA in HIV- 1–infected individuals on suppressive antiretroviral therapy. JCI Insight 5, (2020).

98. Palmer, S. et al. Low-level viremia persists for at least 7 years in patients on suppressive antiretroviral therapy. Proc Natl Acad Sci U S A 105, 3879–84 (2008).

99. Wong, J. K. et al. Reduction of HIV-1 in blood and lymph nodes following potent antiretroviral therapy and the virologic correlates of treatment failure. Proceedings of the National Academy of Sciences 94, 12574–12579 (1997).

100. Yukl, S. A. et al. The Distribution of HIV DNA and RNA in Cell Subsets Differs in Gut and Blood of HIV-Positive Patients on ART: Implications for Viral Persistence. J Infect Dis 208, 1212–1220 (2013).

101. Belmonte, L. et al. The intestinal mucosa as a reservoir of HIV-1 infection after successful HAART. AIDS 21, 2106–2108 (2007).

102. Asowata, O. E., et al. Irreversible depletion of intestinal CD4+ T cells is associated with T cell activation during chronic HIV infection. JCI Insight 6, (2021).

103. Bergler, W., Adam, S., Gross, H. J., Hörmann, K. & Schwartz-Albiez, R. Age-dependent altered proportions in subpopulations of tonsillar lymphocytes. Clin Exp Immunol 116, 9 (1999).

104. Saluzzo, S. et al. Delayed antiretroviral therapy in HIV-infected individuals leads to irreversible depletion of skin- and mucosa-resident memory T cells. Immunity 54, 2842–2858.e5 (2021).

105. Wang, X. et al. Distinct Expression Patterns of CD69 in Mucosal and Systemic Lymphoid Tissues in Primary SIV Infection of Rhesus Macaques. PLoS One 6, e27207 (2011).

106. Reuschl, A.-K. et al. HIV-1 Vpr drives a tissue residency-like phenotype during selective infection of resting memory T cells. Cell Rep 39, 110650 (2022).

107. Banga, R. et al. PD-1+ and follicular helper T cells are responsible for persistent HIV-1 transcription in treated aviremic individuals. Nat Med 22, 754–761 (2016).

108. Tsai, P. et al. In vivo analysis of the effect of panobinostat on cell-associated HIV RNA and DNA levels and latent HIV infection. Retrovirology 13, 36 (2016).

109. Søgaard, O. S. et al. The Depsipeptide Romidepsin Reverses HIV-1 Latency In Vivo. PLoS Pathog 11, e1005142 (2015).

110. Rasmussen, T. A. et al. Panobinostat, a histone deacetylase inhibitor, for latent-virus reactivation in HIV-infected patients on suppressive antiretroviral therapy: a phase 1/2, single group, clinical trial. Lancet HIV 1, e13–e21 (2014).

111. McBrien, J. B. et al. Robust and persistent reactivation of SIV and HIV by N-803 and depletion of CD8+ cells. Nature 578, 154–159 (2020).

112. Jones, R. B. et al. A Subset of Latency-Reversing Agents Expose HIV-Infected Resting CD4+ T-Cells to Recognition by Cytotoxic T-Lymphocytes. PLoS Pathog 12, e1005545 (2016).

113. Miller, J. S. et al. Safety and virologic impact of the IL-15 superagonist N-803 in people living with HIV: a phase 1 trial. Nat Med 28, 392–400 (2022).

114. Chehimi, J. et al. IL-15 enhances immune functions during HIV infection. J Immunol 158, 5978–87 (1997).

115. Garrido, C. et al. Interleukin-15-Stimulated Natural Killer Cells Clear HIV-1- Infected Cells following Latency Reversal *Ex Vivo*. J Virol 92, (2018).

116. Li, Y., Gao, H., Clark, K. M. & Shan, L. IL-15 enhances HIV-1 infection by promoting survival and proliferation of CCR5+CD4+ T cells. JCI Insight 8, (2023).

117. Manganaro, L. et al. IL-15 regulates susceptibility of CD4 ^+^ T cells to HIV infection. Proceedings of the National Academy of Sciences 115, (2018).

118. Schenkel, J. M. et al. IL-15 independent maintenance of tissue resident and boosted effector memory CD8 T cells. doi:10.4049/jimmunol.1502337.

119. Geginat, J., Sallusto, F. & Lanzavecchia, A. Cytokine-driven Proliferation and Differentiation of Human Naive, Central Memory, and Effector Memory CD4+ T Cells. J Exp Med 194, 1711–1720 (2001).

120. Antonucci, J. M. et al. SAMHD1 Impairs HIV-1 Gene Expression and Negatively Modulates Reactivation of Viral Latency in CD4+ T Cells. J Virol 92, (2018).

121. Ruffin, N. et al. Low SAMHD1 expression following T-cell activation and proliferation renders CD4+ T cells susceptible to HIV-1. AIDS 29, 519–530 (2015).

122. Nixon, C. C. et al. Systemic HIV and SIV latency reversal via non-canonical NF-κB signalling in vivo. Nature 578, 160–165 (2020).

123. Mavigner, M. et al. CD8 Lymphocyte Depletion Enhances the Latency Reversal Activity of the SMAC Mimetic AZD5582 in ART-Suppressed Simian Immunodeficiency Virus-Infected Rhesus Macaques. J Virol 95, (2021).

124. Bullen, C. K., Laird, G. M., Durand, C. M., Siliciano, J. D. & Siliciano, R. F. New ex vivo approaches distinguish effective and ineffective single agents for reversing HIV-1 latency in vivo. Nat Med 20, 425–429 (2014).

125. Laird, G. M. et al. Ex vivo analysis identifies effective HIV-1 latency–reversing drug combinations. Journal of Clinical Investigation 125, 1901–1912 (2015).

126. Perelson, A. S., Neumann, A. U., Markowitz, M., Leonard, J. M. & Ho, D. D. HIV- 1 Dynamics in Vivo: Virion Clearance Rate, Infected Cell Life-Span, and Viral Generation Time. Science (1979) 271, 1582–1586 (1996).

127. Wei, X. et al. Viral dynamics in human immunodeficiency virus type 1 infection. Nature 373, 117–122 (1995).

128. Telwatte, S. et al. Gut and blood differ in constitutive blocks to HIV transcription, suggesting tissue-specific differences in the mechanisms that govern HIV latency. PLoS Pathog 14, e1007357 (2018).

129. Elliott, J. H. et al. Activation of HIV Transcription with Short-Course Vorinostat in HIV-Infected Patients on Suppressive Antiretroviral Therapy. PLoS Pathog 10, e1004473 (2014).

130. Lenasi, T., Contreras, X. & Peterlin, B. M. Transcriptional interference antagonizes proviral gene expression to promote HIV latency. Cell Host Microbe 4, 123–33 (2008).

131. Pearson, R. et al. Epigenetic Silencing of Human Immunodeficiency Virus (HIV) Transcription by Formation of Restrictive Chromatin Structures at the Viral Long Terminal Repeat Drives the Progressive Entry of HIV into Latency. J Virol 82, 12291–12303 (2008).

132. Han, Y. et al. Orientation-Dependent Regulation of Integrated HIV-1 Expression by Host Gene Transcriptional Readthrough. Cell Host Microbe 4, 134–146 (2008).

133. Schröder, A. R. W. et al. HIV-1 Integration in the Human Genome Favors Active Genes and Local Hotspots. Cell 110, 521–529 (2002).

134. Ho, Y.-C. et al. Replication-Competent Noninduced Proviruses in the Latent Reservoir Increase Barrier to HIV-1 Cure. Cell 155, 540–551 (2013).

135. Bruner, K. M. et al. A quantitative approach for measuring the reservoir of latent HIV-1 proviruses. Nature 566, 120–125 (2019).

