## Supplemental figures and tables for "Identification of the Inducible HIV reservoir in Tonsillar, Intestinal and Cervical Tissue Models of HIV Latency"

### Supplementary Figure 1

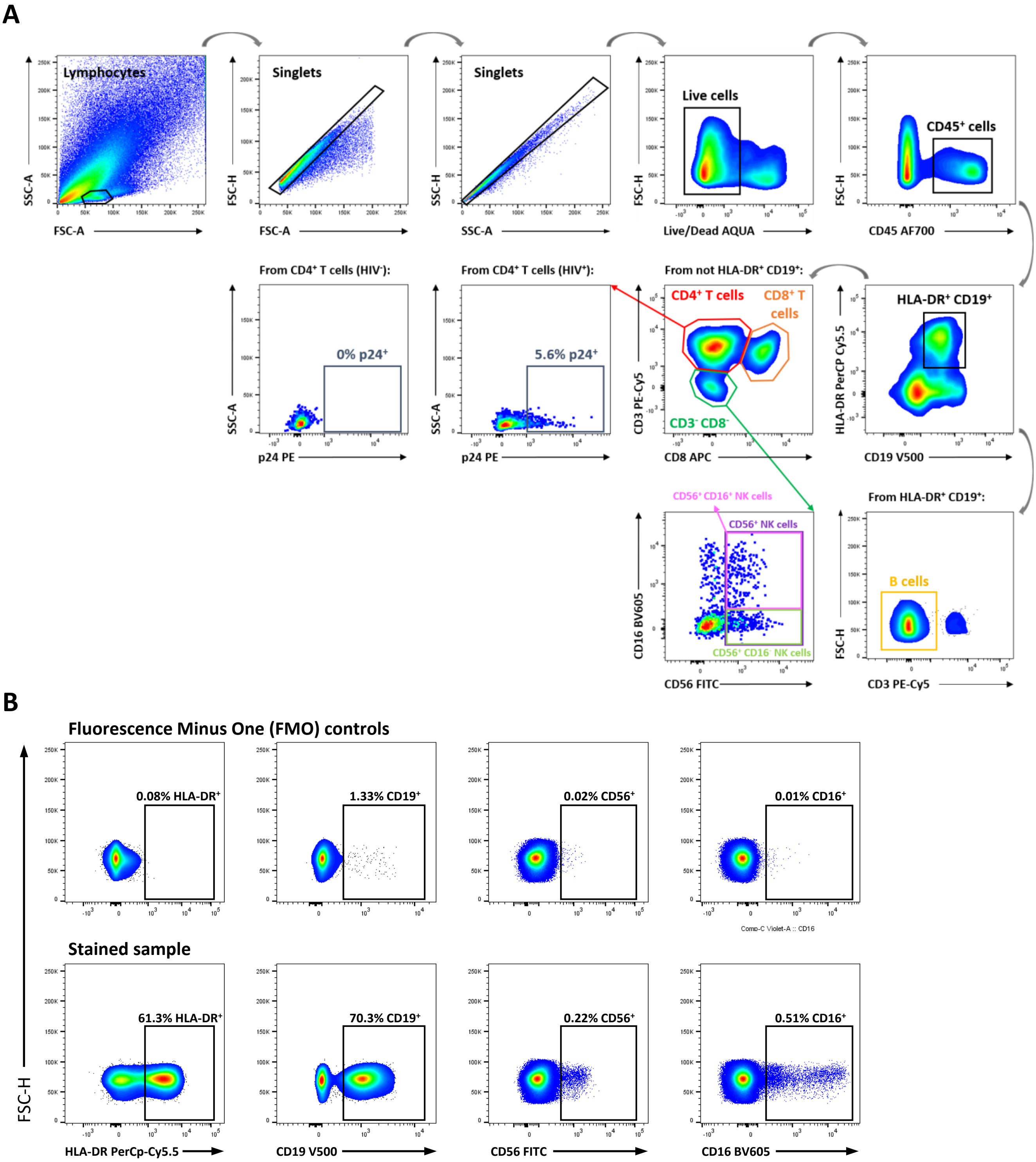

### Supplementary Figure 2

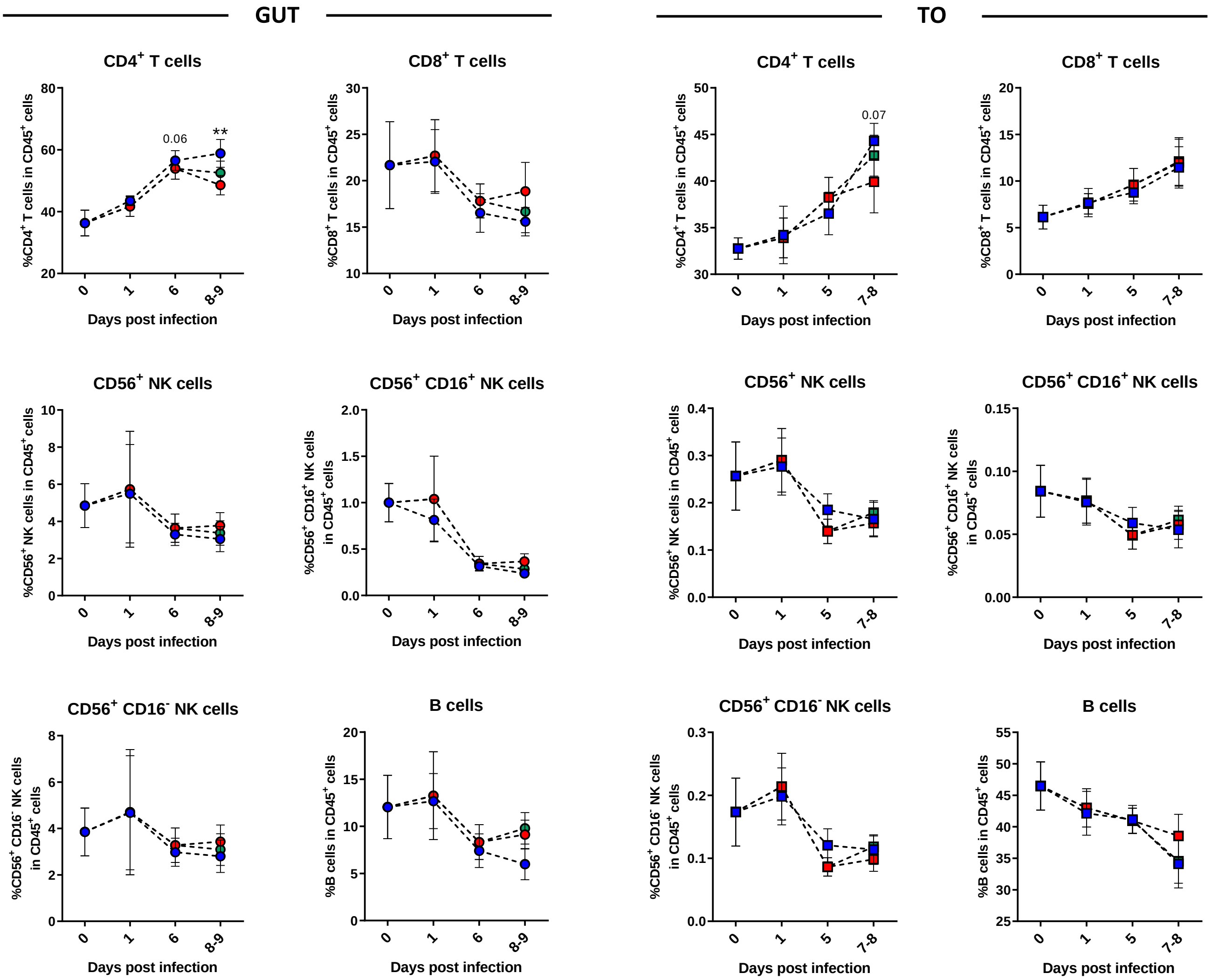

### Supplementary Figure 3

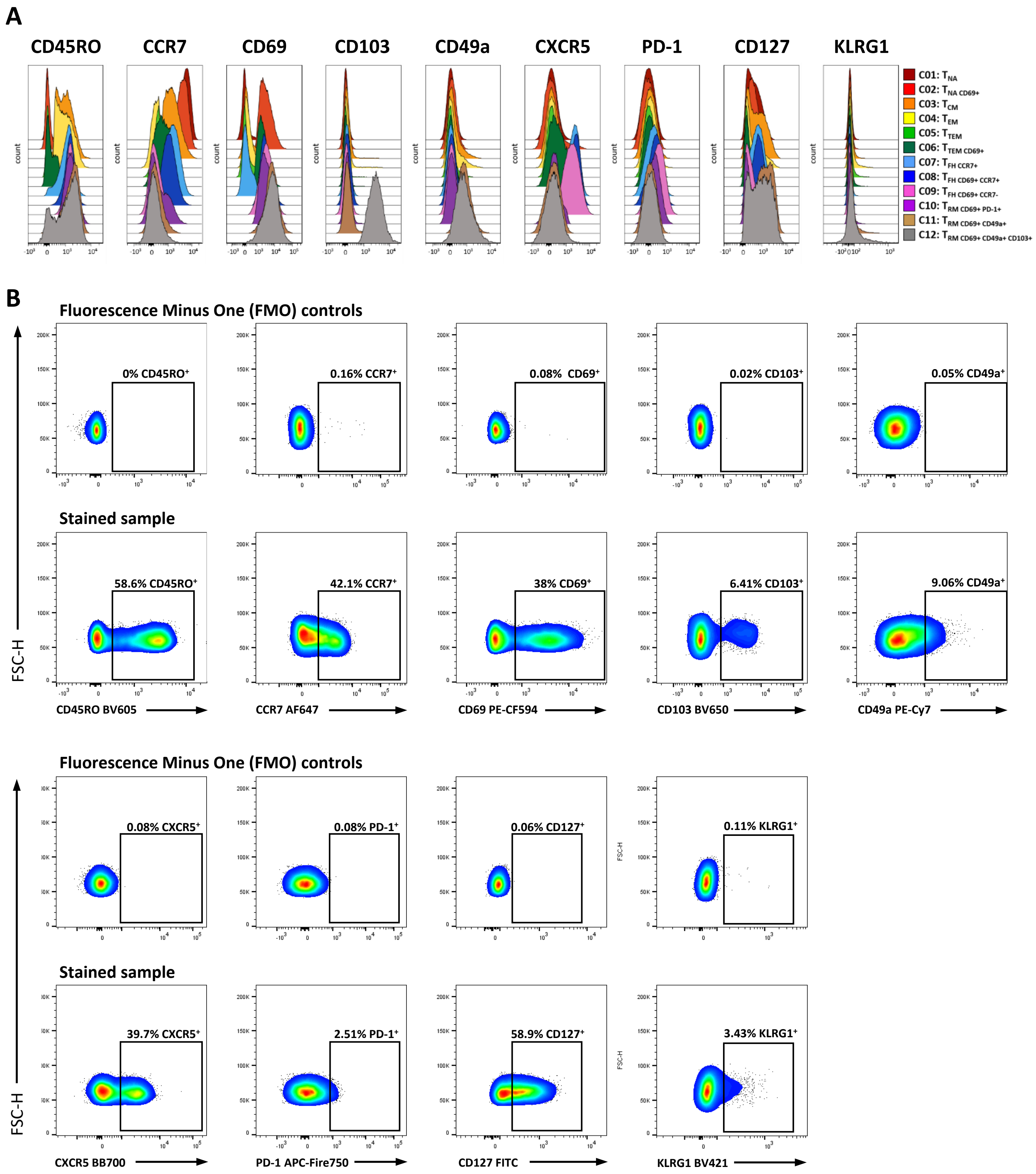

### Supplementary Figure 4

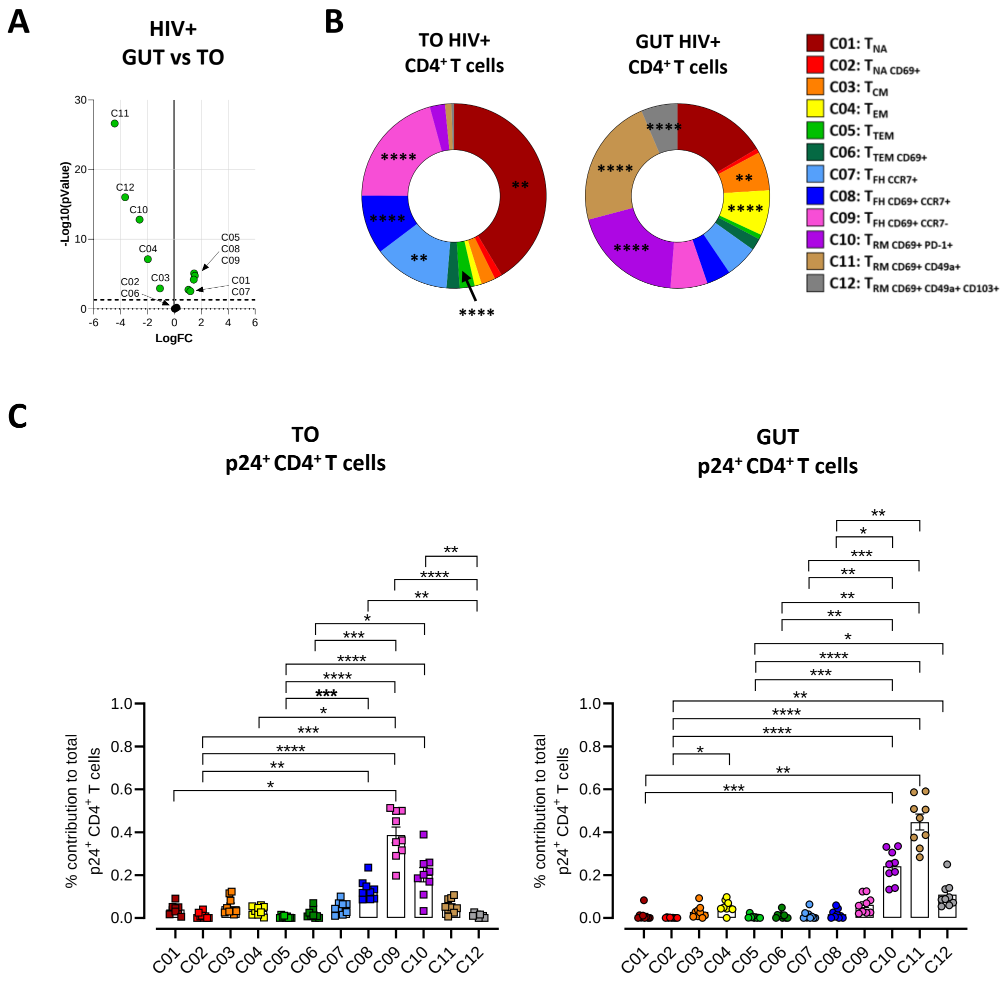

### Supplementary Figure 5

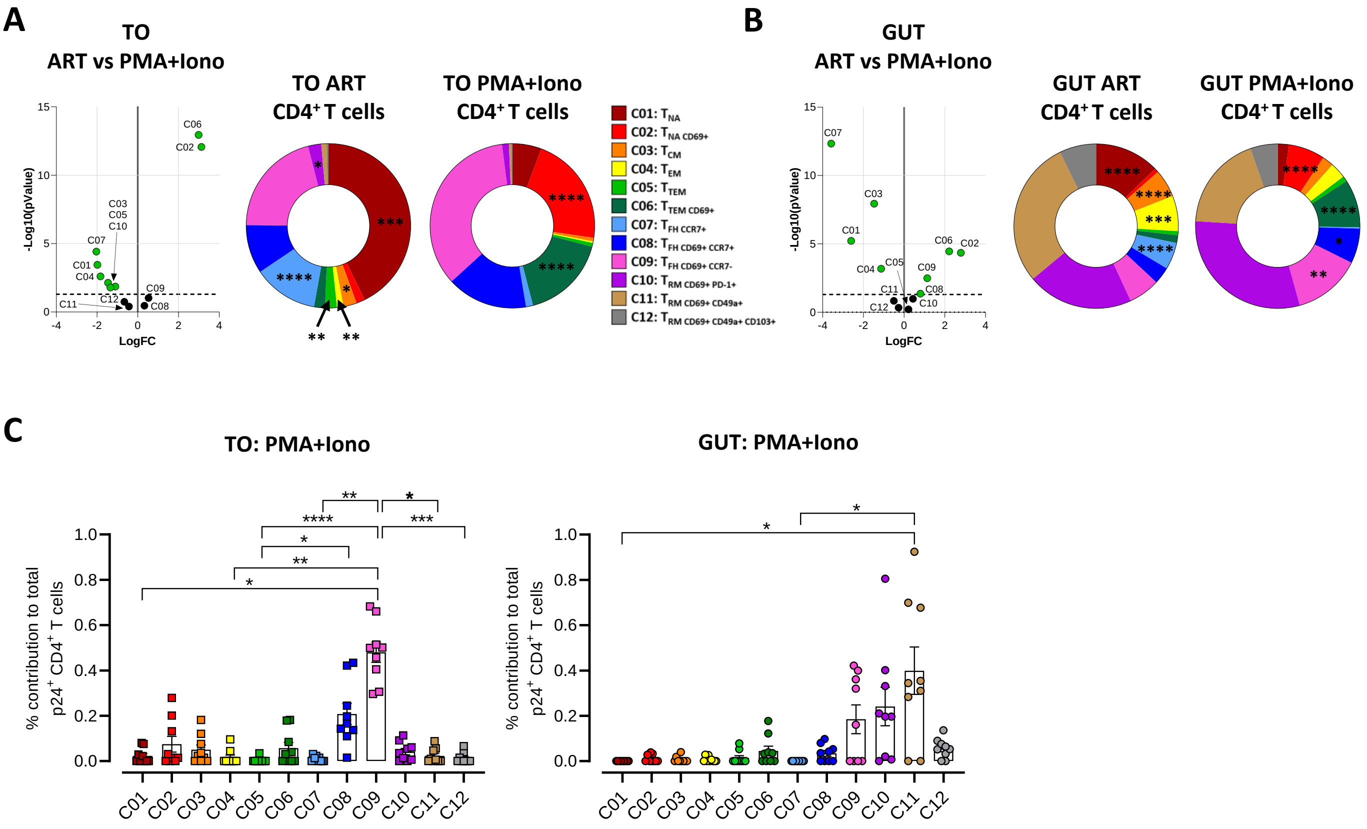

### Supplementary Figure 6

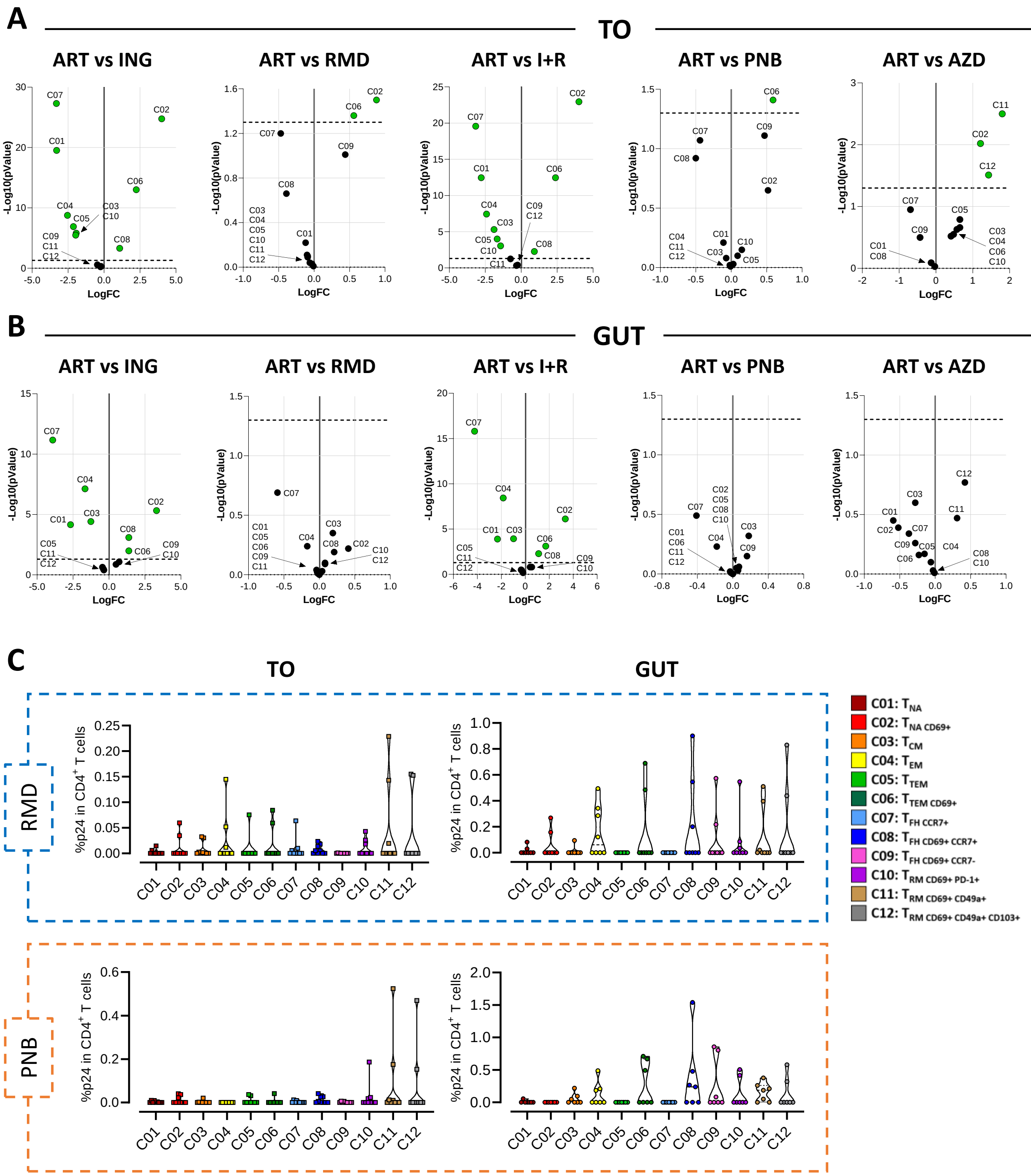

### Supplementary Figure 7

**A**

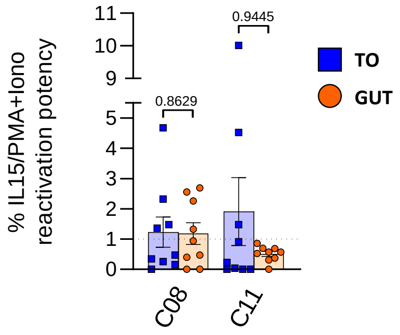

#### Supplementary Table 1

| Tissue | Donors (n) | Gender [no. (%)] | Age median [no. (range)] |
|---|---|---|---|
| Tonsil (TO) | 34 | Male: 20 (59%)<br>Female: 14 (41%) | 5 (2-17) |
| Colon (GUT) | 41 | Male: 23 (56%)<br>Female: 18 (44%) | 71 (37-93) |
| Cervix (CVX) | 45 | Female: 45 (100%) | 55 (35-86) |
